# Small molecule influence on Caudal fin regeneration in Zebrafish: A proteomic based study

**DOI:** 10.1101/2025.03.01.640803

**Authors:** Anusha P V, Shailvi Tewari, Catherine Philip, Anushka Arvind, Ifra Iyoob, Mohammed M Idris

**Author notes:** **Corresponding Author:** Dr. Mohammed M Idris Senior Principal Scientist, CSIR-CCMB, Hyderabad, India.

## Abstract

Dietary and addictive small molecules play a significant role in altering *in vivo* conditions. Due to their minuscule size, these molecules can seamlessly traverse tissues and cellular membranes, influencing key biological processes such as cellular growth, differentiation, and intracellular communication, which are crucial for tissue regeneration. The zebrafish (*Danio rerio*) serves as an excellent model for studying regenerative growth due to its remarkable ability to regrow amputated appendages. In this study, we systematically evaluated the effect of small molecules, including ethanol (0.5%), glucose (1%), and NaCl (0.2%), on zebrafish caudal fin regeneration over a 7-day period. Regenerative growth analysis indicated delayed fin regrowth across all treated groups, with ethanol exposure showing the most significant impairment. Behavioural assessments revealed significant stress-induced locomotor alterations in treated groups, with the ethanol-exposed group exhibiting the most pronounced reduction in total distance moved and velocity. Proteomic profiling using label-free quantification (LFQ) identified 113, 257, and 178 differentially expressed proteins in ethanol, glucose, and NaCl-treated groups, respectively. Subsequent validation using the iTRAQ labeling approach confirmed 16 commonly dysregulated proteins across all conditions, highlighting a shared molecular response associated with stress and repair mechanisms. Pathway enrichment analysis mapped differentially expressed proteins to various canonical signaling pathways, including GP6 signaling, mitochondrial dysfunction, RHO GTPase cycling, antigen processing, and metabolic regulation. Ingenuity Pathway Analysis (IPA) further revealed associations with disease and function networks specific to each treatment condition. Our findings provide valuable insights into how metabolic and ionic perturbations influence zebrafish fin regeneration at the molecular level, offering a deeper understanding of tissue repair mechanisms under stressed conditions.

## Introduction

The Zebrafish (*Danio rerio*) is a fascinating and explorative animal model with a 70% homology to the human reference genome, known for its ability to exhibit an epimorphic regenerative growth in majority of its organs and tissues such as the eyes, heart, spinal cord, fin, etc [1, 2]. The zebrafish is considered an ideal model for studying these mechanisms involved in limb and organ regeneration in vertebrates, with a particular focus on the Caudal fin. The Caudal fin is a non muscularized dermal fold that is stabilized by 16−18 main segmented and occasionally bifurcated rays spanned by soft interlay tissue [3]. Post amputation, it exhibits the process of Epimorphic regeneration consisting of four major events, 1. Injury/Haemostasis, 2. Re-epithelialisation, 3. Blastema formation and 4. Differentiation.

The first stage of the regenerative process consists of wound healing and rapid re-epithelialization, steps that are similar to mammalian tissue regeneration [4]. Following this is the formation of a layer of mesenchymal cells situated on top of the amputated area known as the blastema, after which it transitions to a regenerative outgrowth via the differentiation process [1]. Zebrafish Caudal fin regeneration comprises the following four stages: “epithelialization or wound healing” (0–1-day post-amputation (dpa)), “blastema formation” (1–2dpa), “regenerative outgrowth” (2–7dpa), and “termination”. At 1dpa, a drift of the proximal epidermis is seen that covers the amputated region, followed by inflammation. Histolysis, cell dedifferentiation and blastema formation start at 2dpa. Subsequently, between 3-7dpa, regenerative outgrowth is seen with the robust proliferation of dedifferentiated cells, progressive redifferentiation, and morphogenesis take place. After 8-10dpa, the fin regenerates completely and regenerative homeostasis is achieved [5].

Small molecules are classified as such due to their small size (< 900 Da) and are known to penetrate various tissues and cells via passive or facilitated diffusion and through ion channels; these molecules have pharmacological significance, i.e., in precision medicine and targeted therapeutic drugs. A few known small molecules like ethanol, glucose, and sodium chloride can interact in various ways, including affecting glucose transport, electrolyte concentrations, and fluid balance in the living body including the regeneration process. It is pertinent to mention here that each of these molecules has limitations regarding the beneficial effect on various organs. For example, ethanol can increase the energy state by stimulating dopaminergic neural activity signalling reward in the brain, whereas chronic ethanol metabolism results in fatty liver and general metabolic dysfunction in the liver [6]. Similarly, sodium chloride is essential for the support of various body functions such as the conduction of nerve impulses, contraction and relaxation of muscles, and maintenance of the proper balance of water and minerals. However, when consumed in excess, it causes increased blood pressure, which can lead to heart disease, stroke, and premature death [7]. Similarly, glucose is a diverse monosaccharide that has isometric forms such as galactose and fructose, lactose and sucrose (disaccharides), or starch (polysaccharide) upon its entry into the human body, where it is mostly converted to ATP and stored as Glycogen [8] for future use. Acute and severe reduction of brain glucose can lead to impairment of cognitive and reflex function, autonomic failure, seizures, loss of consciousness, and permanent and irreversible brain damage [9].

With the use of zebrafish as a tool, researchers have shown that it is a good model of Foetal Alcohol Spectrum Disorders (FASD) [10]. Additionally, in zebrafish larvae, the hypothalamic-pituitary-adrenal (HPA) axis activity was found to exhibit increased and disrupted neurotransmitter metabolism due to ethanol exposure [11]. Likewise, excess sodium chloride exposure led to reduced sleep in larval zebrafish, leading to increased cortisol levels [12]. Considering how past studies have utilized zebrafish embryos [13, 14, 15] as well as adult fish [15, 16] it appears a promising model to study these various pathways affected by high-glucose exposure like mammals[16].Reflecting on previous literature within this discipline, it would be in the best interest to know how these molecules individually affect the regeneration process. This present study is designed as one of the first to evaluate the individual effects of these 3 molecules (Ethanol, Sodium Chloride, and Glucose) on the caudal fin epimorphic regeneration process.

## Materials and Methodology

### Animal Housing and Regeneration experiment

Adult wild-type zebrafish (6-12 months old) were collected from the CCMB zebrafish laboratory and maintained on a 14-hour light/10-hour dark cycle. Water temperature and pH are maintained under standard laboratory conditions. The zebrafish were housed in four different conditioned tanks with normal water as Control, 0.5% Ethanol, 0.2% NaCl and 1% Glucos-treated water for the 7-day time period. The zebrafish were anesthetized with 0.1% tricaine and amputated on the caudal fin distal region for regeneration experiments [5]. The regenerated caudal fins were collected at 0hpa, 1dpa, 2dpa, and 7day post amputation. The experiment protocol was approved by the Centre for Cellular and Molecular Biology institutional animal ethics committee (IAEC/CCMB/Protocol #50/2013).

### Behavioural Analysis

The novel tank test was performed to analyse the behaviours of the control and treatment groups simultaneously upon amputation and non-amputation for a period of 7 days. The novel tank test (NTT) mimics the open field and elevated plus maze tests used to assess anxiety [17]. After an acclimatization period of 5 min, zebrafish were placed individually in a narrow 15×12×25-cm tank with a water depth of 15 cm divided into three equal, virtually horizontal sections and demarcated by a line on the outside of the tank wall. In the 2-min novel tank test, the time spent by the fish in the different levels of the tank (bottom, middle or upper level) was measured to assess the level of stress. A preference for the bottom two levels, less frequent venturing into the upper level of the tank and a lower number of crosses in the swim area is suggestive of increased stress or anxiety. The analysis was performed using Ethiovision software.

### Proteomic Analysis

The total protein of all the selected time points of regenerating caudal fin tissues of each group was extracted [18] and quantified using Amido black assay with bovine serum albumin as standard [5,18,19]. Both Label-free Quantification as well as Isobaric Taq for Relative and Absolute Quantification (iTRAQ) based quantitative proteomics were carried out for all the time points of regeneration against control (0hpa).

### Label-free Quantitative Proteomics

100 µg of total protein from each experimental group with 1dpa, 2dpa and 7dpa tissue lysed protein samples underwent electrophoresis on 10% SDS-PAGE gels, which were then stained with Commassie R250, destained, and the gel excised into four fractions based on molecular weight. In-gel digestion using trypsin was followed by purification of the digested peptides using C-18 spin columns (Thermo Scientific). The purified peptides were reconstituted in 5% acetonitrile (ACN) and 0.2% formic acid before undergoing Liquid Chromatography Mass Spectrometry (LCMS/MSMS) analysis using an Orbitrap Velos Nano analyzer (Q-Exactive HF). Proteomic data obtained from the mass spectrometer were analyzed against *Danio rerio*proteome database. Differential expression in proteins was estimated relative to the non-treated control negative samples alongside the 0hpa tissue control for regeneration.

### iTRAQ Label-based Quantitative Proteomics

Isobaric Taq for Relative and Absolute Quantification (iTRAQ) based quantitative proteomics was carried out for all the selected time points of regeneration (1dpa, 2dpa and 7dpa) for each of the treatment groups against the control (non-treated) group in succession to a trypsin based 10% SDS gel digestion. All the digested peptides were labelled with iTRAQ 4-plex. iTRAQ labels were designated as follows: 114 for non-treated Control, 115 for ethanol-treated, 116 for NaCl-treated, and 117 for glucose-treated peptides. All the labeled peptides were pooled according to the fractions and purified using C-18 spin columns (Thermo Scientific). Labelled and purified peptides were analyzed using the LC-MS/MS Orbitrap Velos Nano analyser (Thermo Scientific). The resulting raw data was analyzed using 1% FDR percolator and XCorr (Score Vs Charge) and D. rerio database. Proteins having more than one log-fold change in the selected time points were recruited for the study.

### Data Submission, Heatmap and Network Pathway Analysis

The Proteomics data generated from the study were deposited to the ProteomeXchange consortium via the PRIDE partner repository with the dataset Accession No. PXD060144. The differentially expressed genes/proteins were analyzed for the heat map and network pathway analysis involving the heatmapper portal (www.heatmapper.ca) and Ingenuity pathway analysis (IPA) software respectively. The heat maps were generated for the differentially expressed genes and proteins involving hierarchical cluster analysis. Network pathways were generated using IPA software for the canonical pathway as well as disease and functions.

## Results

### Effect of Ethanol, Glucose and NaCl in Eegeneration

Based on our study we found that adult zebrafish can tolerate 0.5% Ethanol, 1% Glucose and 0.2% NaCl concentration without having any lethal effect (LD50). Exposure of 0.5% Ethanol, 1% Glucose and 0.2 % NaCl concentration was found to affect the regrowth of the caudal fin in zebrafish upon amputation (figure 1a). The rate of regeneration was found compromised in both the lobe and cleft part of the fin tissue at the end of 7dpa, as previously described [5]. Upon measurement of the regenerated fin, the simultaneous regeneration and treatment period of 7days showed variation among Glucose and NaCl-treated groups whereas the Ethanol-treated group showed significant variation of growth in both lobe and cleft compared to the Control group (figure 1b).

**Figure 1:**
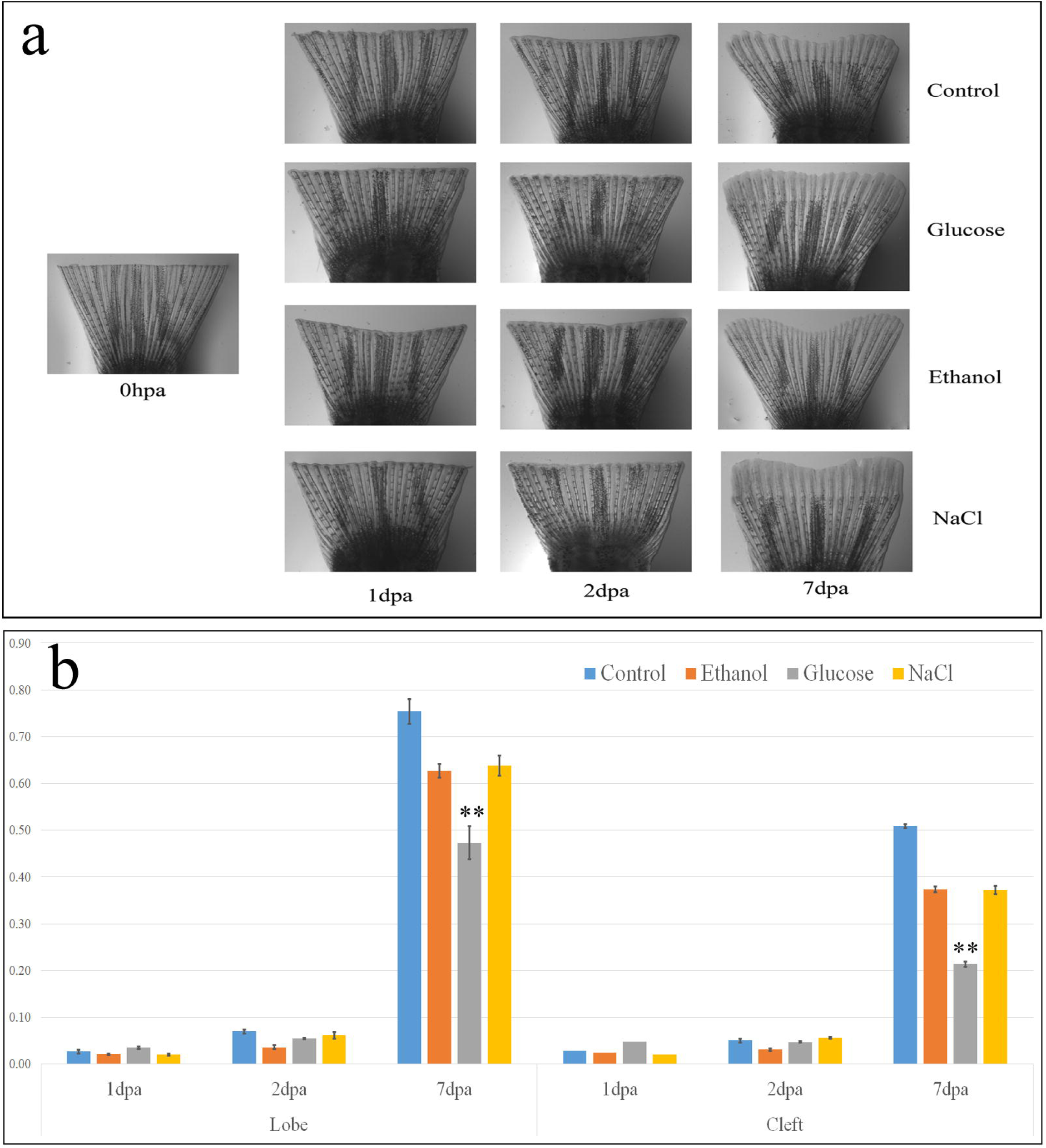
Regeneration of zebrafish caudal fins post amputation at different stages under different treatment conditions - (a). Amputated zebrafish fin at 0hpa and the subsequent images depicting regenerating fins at 1, 2, and 7 days post-amputation (dpa) in control, glucose, ethanol, and NaCl-treated groups, (b). Quantification of regenerative growth in the lobe and cleft regions across different time points (1, 2, and 7 dpa) for all treatment groups. Statistical significance is indicated (p < 0.01).

Based on our behavioural analysis study, we found that exposure to 0.5% Ethanol, 1% Glucose and 0.2% NaCl induced alterations in zebrafish behaviour that reflected stressed conditions upon the treatment under parameters such as distance and velocity (figure 2). The total distance moved (figure 2a) and velocity (figure 2b) were found to be decreased in the treated group compared to non-treated control group. There were also more alterations among the treated group compared to the group subjected to simultaneous amputation and treatment. Moreover, we found that the movement of ethanol-treated fish was significantly lower than that of the control fish. In the treated groups, the fish were found to have spent more time in the bottom levels of the tank than in the upper level.

**Figure 2:**
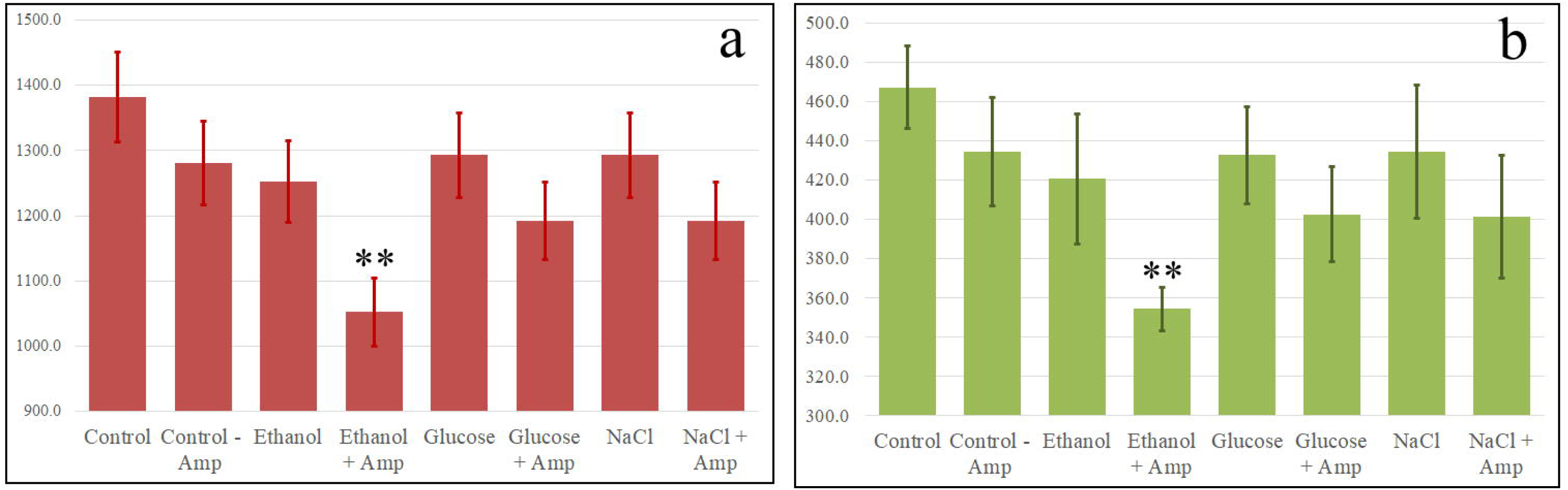
Effect of small molecule exposure on zebrafish locomotor behavior. (a) Total distance moved by zebrafish under different treatment conditions, including control, ethanol (0.5%), glucose (1%), and NaCl (0.2%), with and without amputation (p<0.01). (b) Average velocity of zebrafish across different treatment groups (p < 0.01).

Based on our proteomic label-free quantification analysis, we identified 113, 257, and 178 differentially expressed proteins in zebrafish caudal fin tissues following exposure to 0.5% ethanol, 1% glucose, and 0.2% NaCl, respectively (Tables 1,2 & 3). A total of 16 proteins were found to be differentially expressed across all three exposure groups, validated using the iTRAQ-labeled quantification method (Table 4). These findings highlight a shared set of molecular responses to these treatments, potentially linked to common stress or repair mechanisms.

**Table 1:** List of Proteins found differentially expressed in zebrafish caudal fin tissue upon exposure to Ethanol.

| S.No | Accession | Description | Coverage [%] | # Peptides | # PSMs | 1dpa | 2dpa | 7dpa |
| --- | --- | --- | --- | --- | --- | --- | --- | --- |
| 1 | XP_005163812.1 | brain-specific angiogenesis inhibitor 1-associated protein 2 isoform X1 | 29 | 10 | 35 | -1.5 | 0.2 | -0.6 |
| 2 | NP_001002343.1 | guanylate-binding protein 1 | 21 | 9 | 40 | -1.5 | -0.7 | -0.1 |
| 3 | NP_001032197.2 | uncharacterized protein LOC641325 precursor | 45 | 11 | 160 | -1.2 | 0.3 | 0.2 |
| 4 | NP_001002645.3 | enoyl-CoA delta isomerase 2, mitochondrial | 16 | 4 | 20 | -1.2 | -0.8 | 0.3 |
| 5 | NP_957202.1 | tubulin gamma-1 chain isoform 1 | 11 | 3 | 6 | -1.2 | 0.1 | 0.6 |
| 6 | NP_570982.2 | tenascin isoform 1 precursor | 20 | 21 | 120 | -1.2 | -0.1 | -0.2 |
| 7 | NP_958472.2 | ATP-binding cassette sub-family F member 2a | 10 | 5 | 10 | -1.2 | 0.1 | -0.3 |
| 8 | XP_009296840.2 | E3 ubiquitin-protein ligase RNF31 | 3 | 2 | 2 | -1.2 | 2.0 | 0.1 |
| 9 | NP_991281.3 | GDP-fucose protein O-fucosyltransferase 1 precursor | 14 | 4 | 18 | -1.1 | 0.2 | -0.1 |
| 10 | XP_005169569.1 | protein phosphatase 1G isoform X1 | 2 | 1 | 2 | -1.1 | -0.1 | 0.3 |
| 11 | XP_005155921.1 | matrilin-4 isoform X1 | 6 | 4 | 6 | -1.1 | -0.8 | -0.7 |
| 12 | NP_997956.1 | adenosine kinase isoform 1 | 16 | 5 | 13 | -1.1 | -0.9 | -1.3 |
| 13 | NP_001017626.1 | prenylcysteine oxidase 1 precursor | 12 | 5 | 10 | -1.0 | 1.4 | 0.3 |
| 14 | NP_001278325.1 | parafibromin isoform 1 | 8 | 3 | 11 | -1.0 | -0.2 | 0.1 |
| 15 | NP_775339.1 | 40S ribosomal protein S5 | 31 | 5 | 58 | -1.0 | -0.2 | -0.3 |
| 16 | NP_001002715.1 | uncharacterized protein LOC436988 | 12 | 4 | 17 | -0.7 | -1.0 | 0.5 |
| 17 | XP_005160712.1 | vacuolar protein sorting-associated protein VTA1 homolog isoform X1 | 21 | 3 | 4 | -0.7 | -1.0 | -0.6 |
| 18 | NP_001315090.1 | dolichyl-diphosphooligosaccharide--protein glycosyltransferase subunit STT3B | 8 | 6 | 26 | -0.7 | 0.4 | -1.3 |
| 19 | NP_001038237.1 | leucine-rich alpha-2-glycoprotein precursor | 34 | 11 | 49 | -0.6 | 0.2 | 1.0 |
| 20 | NP_955365.1 | pyruvate kinase PKM | 48 | 24 | 333 | -0.5 | -0.5 | 1.2 |
| 21 | NP_957366.1 | reticulon-3 isoform 1 | 22 | 6 | 30 | -0.5 | 0.7 | -1.0 |
| 22 | NP_001009989.1 | protein kinase, cAMP-dependent, regulatory, type I, alpha (tissue specific extinguisher 1) a | 27 | 8 | 66 | -0.5 | -0.6 | 1.0 |
| 23 | XP_017207875.1 | ectonucleotide pyrophosphatase/phosphodiesterase family member 2-like | 14 | 9 | 42 | -0.4 | 1.0 | 0.8 |
| 24 | XP_002665524.2 | carcinoembryonic antigen-related cell adhesion molecule 5 isoform X1 | 16 | 6 | 85 | -0.4 | 0.1 | -1.3 |
| 25 | AAI55791.1 | LOC555531 protein, partial | 6 | 2 | 8 | -0.4 | -0.1 | -1.0 |
| 26 | NP_001122009.1 | host cell factor 1b | 16 | 18 | 77 | -0.4 | 0.4 | -1.0 |
| 27 | NP_571340.1 | vitellogenin 3, phosvitinless precursor | 31 | 24 | 104 | -0.4 | 0.2 | 1.2 |
| 28 | NP_997787.2 | alpha-2-HS-glycoprotein 1 precursor | 14 | 5 | 37 | -0.4 | 0.1 | -1.6 |
| 29 | XP_694950.3 | protein-glutamine gamma-glutamyltransferase K | 13 | 8 | 28 | -0.3 | 0.3 | -1.0 |
| 30 | NP_944595.2 | DNA replication licensing factor MCM4 | 30 | 18 | 118 | -0.3 | 0.3 | -1.5 |
| 31 | XP_001335151.4 | neuroblast differentiation-associated protein AHNAK-like isoform X1 | 3 | 8 | 27 | -0.2 | 0.0 | 1.4 |
| 32 | NP_998255.2 | scinderin like b | 52 | 36 | 1088 | -0.2 | 0.0 | -1.1 |
| 33 | NP_571765.1 | sodium/potassium-transporting ATPase subunit alpha-1b | 26 | 19 | 290 | -0.2 | 0.0 | -1.5 |
| 34 | NP_956468.2 | uridine 5'-monophosphate synthase | 22 | 9 | 29 | -0.2 | -1.0 | 0.7 |
| 35 | NP_001003412.1 | clathrin interactor 1a | 9 | 4 | 16 | -0.2 | -0.5 | 1.4 |
| 36 | XP_017212305.1 | AP-1 complex subunit gamma-1 isoform X1 | 16 | 10 | 25 | -0.1 | -1.1 | 0.4 |
| 37 | NP_001071636.2 | sulfotransferase family 2, cytosolic sulfotransferase 3 | 77 | 25 | 290 | -0.1 | -0.4 | -1.5 |
| 38 | XP_021336506.1 | calcium/calmodulin-dependent protein kinase type II subunit gamma isoform X44 | 9 | 4 | 37 | -0.1 | 0.0 | -1.4 |
| 39 | NP_001004539.1 | pigment epithelium-derived factor precursor | 35 | 8 | 37 | -0.1 | -1.0 | 0.4 |
| 40 | XP_696031.6 | laminin subunit alpha-3 | 22 | 21 | 107 | -0.1 | -0.1 | -1.0 |
| 41 | XP_017214411.1 | exportin-5 | 2 | 2 | 7 | -0.1 | 0.7 | -1.9 |
| 42 | NP_001108387.1 | C-reactive protein 3 precursor | 24 | 4 | 16 | -0.1 | -0.8 | -1.2 |
| 43 | XP_005173887.2 | desmoplakin-like | 40 | 12 | 156 | -0.1 | -0.2 | -1.2 |
| 44 | XP_017209214.1 | cadherin-like protein 26 | 25 | 16 | 96 | 0.0 | -0.3 | -1.6 |
| 45 | NP_571385.2 | heat shock protein HSP 90-beta | 53 | 51 | 851 | 0.0 | 0.2 | -1.2 |
| 46 | NP_958482.1 | major vault protein | 62 | 50 | 667 | 0.0 | 0.4 | -1.1 |
| 47 | NP_835232.2 | scinderin like a | 50 | 40 | 803 | 0.0 | 0.1 | -1.2 |
| 48 | XP_005163413.1 | phospholipase C, delta 1b isoform X1 | 40 | 29 | 213 | 0.0 | 0.7 | -1.6 |
| 49 | NP_954684.1 | collagen, type I, alpha 1a precursor | 18 | 18 | 185 | 0.1 | 0.2 | -1.4 |
| 50 | NP_001304681.1 | vinculin b | 52 | 51 | 630 | 0.1 | 0.4 | -1.2 |
| 51 | XP_021323922.1 | CUB domain-containing protein 1-like | 8 | 6 | 14 | 0.1 | -0.4 | -1.5 |
| 52 | XP_009303829.1 | fibronectin isoform X2 | 41 | 63 | 844 | 0.1 | -0.2 | -1.3 |
| 53 | XP_683754.2 | RNA-binding protein NOB1 isoform X1 | 9 | 2 | 10 | 0.1 | -0.2 | -2.2 |
| 54 | NP_998219.1 | fibrinogen gamma chain precursor | 42 | 21 | 342 | 0.1 | 0.3 | -1.0 |
| 55 | XP_005164160.1 | periplakin isoform X1 | 54 | 111 | 943 | 0.1 | 0.0 | -1.0 |
| 56 | NP_571761.1 | sodium/potassium-transporting ATPase subunit alpha-1a.1 | 38 | 29 | 439 | 0.1 | 0.2 | -1.4 |
| 57 | XP_021322510.1 | plectin isoform X22 | 55 | 273 | 3023 | 0.1 | 0.0 | -1.3 |
| 58 | NP_001007362.1 | protein SGT1 homolog | 12 | 4 | 12 | 0.1 | 0.3 | -1.1 |
| 59 | XP_021330653.1 | myotrophin isoform X1 | 20 | 3 | 43 | 0.2 | -1.4 | -0.9 |
| 60 | XP_005160999.1 | alpha-adducin isoform X5 | 17 | 9 | 39 | 0.2 | -0.1 | -1.3 |
| 61 | XP_017210833.1 | protein NLRC3 | 20 | 10 | 46 | 0.2 | 0.5 | -1.0 |
| 62 | Q566Y8.1 | RecName: Full=Protein mago nashi homolog | 55 | 6 | 60 | 0.2 | 1.0 | -0.2 |
| 63 | NP_001030159.1 | integrin beta-1b precursor | 16 | 7 | 73 | 0.2 | 0.2 | -1.1 |
| 64 | XP_009289535.1 | neutral alpha-glucosidase AB | 39 | 22 | 130 | 0.2 | 0.3 | -1.1 |
| 65 | NP_998668.1 | splicing factor 3B subunit 3 | 29 | 23 | 108 | 0.2 | 0.2 | -1.3 |
| 66 | XP_005156033.1 | collagen, type I, alpha 1b isoform X1 | 14 | 18 | 275 | 0.2 | -0.1 | -1.6 |
| 67 | NP_001315319.1 | 10 kDa heat shock protein, mitochondrial isoform 1 | 45 | 5 | 72 | 0.2 | -1.0 | 0.3 |
| 68 | XP_021336562.1 | uncharacterized protein LOC563946 isoform X1 | 73 | 39 | 600 | 0.2 | 1.3 | -0.3 |
| 69 | XP_001919291.4 | uncharacterized protein LOC563036 | 38 | 16 | 97 | 0.3 | 0.0 | -1.0 |
| 70 | NP_001002215.2 | microsomal glutathione S-transferase 1.2 | 15 | 3 | 9 | 0.3 | 0.7 | -2.3 |
| 71 | NP_001139027.1 | ribosome-binding protein 1a | 31 | 35 | 120 | 0.3 | 0.0 | -1.1 |
| 72 | NP_958866.1 | dolichyl-diphosphooligosaccharide--protein glycosyltransferase subunit STT3A | 18 | 11 | 62 | 0.3 | -0.3 | -1.0 |
| 73 | NP_892013.2 | collagen alpha-2(I) chain precursor | 18 | 21 | 436 | 0.3 | 0.0 | -1.9 |
| 74 | XP_009304310.1 | host cell factor C1a isoform X1 | 12 | 13 | 58 | 0.3 | 0.2 | -1.4 |
| 75 | NP_001303659.1 | [F-actin]-monooxygenase mical1 | 3 | 2 | 11 | 0.4 | 0.5 | 1.0 |
| 76 | XP_002663145.3 | epidermal growth factor receptor substrate 15 isoform X1 | 6 | 5 | 14 | 0.5 | 0.0 | -1.0 |
| 77 | NP_775333.1 | thrombospondin-4-B precursor | 6 | 4 | 10 | 0.5 | 0.1 | -1.7 |
| 78 | AFS66692.1 | mitochondrial calcium uniporter | 26 | 7 | 52 | 0.5 | 0.3 | -1.4 |
| 79 | NP_997975.2 | structural maintenance of chromosomes 1A, like | 24 | 24 | 81 | 0.5 | 0.3 | -1.4 |
| 80 | XP_021324618.1 | uncharacterized protein LOC110438215 | 29 | 10 | 31 | 0.5 | 0.4 | -1.0 |
| 81 | XP_009296304.1 | plakophilin-3 isoform X1 | 39 | 30 | 325 | 0.6 | 0.1 | 1.4 |
| 82 | AAI65624.1 | F10 protein | 6 | 2 | 14 | 0.6 | 0.0 | -1.1 |
| 83 | XP_005155352.1 | exportin-7 isoform X1 | 7 | 5 | 26 | 0.6 | -1.2 | 1.0 |
| 84 | XP_005165182.1 | apoptosis-inducing factor 1, mitochondrial isoform X1 | 29 | 10 | 77 | 0.6 | 0.6 | -1.3 |
| 85 | XP_005173061.1 | protein transport protein Sec31A isoform X2 | 7 | 7 | 15 | 0.6 | 1.2 | -0.9 |
| 86 | XP_686012.3 | integrin beta-2 | 16 | 7 | 30 | 0.7 | 0.0 | -1.4 |
| 87 | XP_001343636.1 | rho GTPase-activating protein 35 | 4 | 3 | 9 | 0.7 | -0.4 | -1.3 |
| 88 | XP_697241.4 | collagen alpha-1(XVII) chain isoform X1 | 5 | 5 | 14 | 0.7 | 0.0 | -1.5 |
| 89 | XP_021324673.1 | collagen alpha-1(V) chain isoform X1 | 9 | 9 | 35 | 0.7 | 0.5 | -1.4 |
| 90 | AAO61483.1 | fetuin-A | 16 | 6 | 36 | 0.7 | 1.0 | 0.3 |
| 91 | XP_017211203.1 | NACHT, LRR and PYD domains-containing protein 12-like | 24 | 17 | 165 | 0.7 | 0.3 | -1.1 |
| 92 | NP_001073381.1 | reticulon-4a isoform 4-l | 35 | 5 | 28 | 0.8 | 0.3 | -2.1 |
| 93 | XP_017209369.1 | collagen alpha-1(XXI) chain-like isoform X1 | 13 | 9 | 30 | 0.8 | -0.2 | -1.0 |
| 94 | Q7ZUG5.1 | RecName: Full=40S ribosomal protein S21 | 42 | 5 | 112 | 0.8 | -1.1 | -0.2 |
| 95 | XP_017214412.1 | LOW QUALITY PROTEIN: echinoderm microtubule-associated protein-like 4 | 19 | 12 | 37 | 1.0 | 0.3 | -0.7 |
| 96 | AGL92229.1 | MHC class I antigen UIA | 39 | 8 | 60 | 1.0 | -0.3 | 0.2 |
| 97 | Q6DHU1.1 | RecName: Full=Cleft lip and palate transmembrane protein 1-like protein; Short=CLPTM1-like protein | 3 | 1 | 6 | 1.0 | 0.2 | 0.6 |
| 98 | XP_021327607.1 | vacuolar protein sorting-associated protein 13C isoform X1 | 3 | 7 | 18 | 1.0 | 0.3 | 0.7 |
| 99 | AAH96883.1 | Si:dkey-276i5.1 protein, partial | 29 | 19 | 167 | 1.0 | 0.2 | 0.0 |
| 100 | NP_001265725.1 | aldehyde oxidase 5 | 3 | 2 | 4 | 1.1 | 0.2 | 0.4 |
| 101 | XP_002663865.1 | prolyl 3-hydroxylase 1 | 24 | 14 | 35 | 1.1 | 0.3 | -0.6 |
| 102 | NP_001290058.1 | ATP synthase subunit f, mitochondrial | 12 | 1 | 3 | 1.1 | -0.2 | -1.1 |
| 103 | NP_955463.2 | ribosome-binding protein 1b | 13 | 8 | 24 | 1.1 | 0.0 | -2.2 |
| 104 | XP_021337011.1 | alpha-actinin-4 isoform X3 | 67 | 67 | 1275 | 1.1 | 0.9 | -1.5 |
| 105 | NP_001035455.1 | advillin | 11 | 6 | 14 | 1.1 | 1.1 | 0.5 |
| 106 | Q7ZVZ6.1 | RecName: Full=Presequence protease, mitochondrial;<br>AltName: Full=Pitriylsin metalloproteinase 1; Flags:<br>Precursor | 10 | 6 | 28 | 1.2 | 0.1 | 0.4 |
| 107 | XP_001340860.1 | probable ATP-dependent RNA helicase ddx6 | 39 | 13 | 78 | 1.3 | 0.0 | -0.7 |
| 108 | AOA0R4IBK5.1 | RecName: Full=E3 ubiquitin-protein ligase rnf213-alpha;<br>AltName: Full=Mysterin-A; AltName: Full=Mysterin-alpha;<br>AltName: Full=RING finger protein 213-A; AltName:<br>Full=RING finger protein 213-alpha; AltName: Full=RING-type E3 ubiquitin transferase rnf213-alpha | 3 | 9 | 27 | 1.3 | 0.2 | 0.8 |
| 109 | Q6NWE0.1 | RecName: Full=Ester hydrolase C11orf54 homolog | 28 | 5 | 19 | 1.4 | -1.3 | -1.5 |
| 110 | XP_021331113.1 | protein NLRC3-like isoform X2 | 6 | 4 | 53 | 1.5 | 0.4 | -1.5 |
| 111 | NP_001338766.1 | eosinophil peroxidase isoform 1 precursor | 8 | 4 | 10 | 1.5 | 0.3 | -1.2 |
| 112 | XP_021323351.1 | enhancer of mRNA-decapping protein 4 isoform X1 | 8 | 6 | 25 | 1.6 | 0.0 | -0.2 |
| 113 | XP_021334459.1 | fer-1-like protein 6 | 1 | 1 | 12 | 2.0 | -0.2 | -0.1 |

**Table 2:**
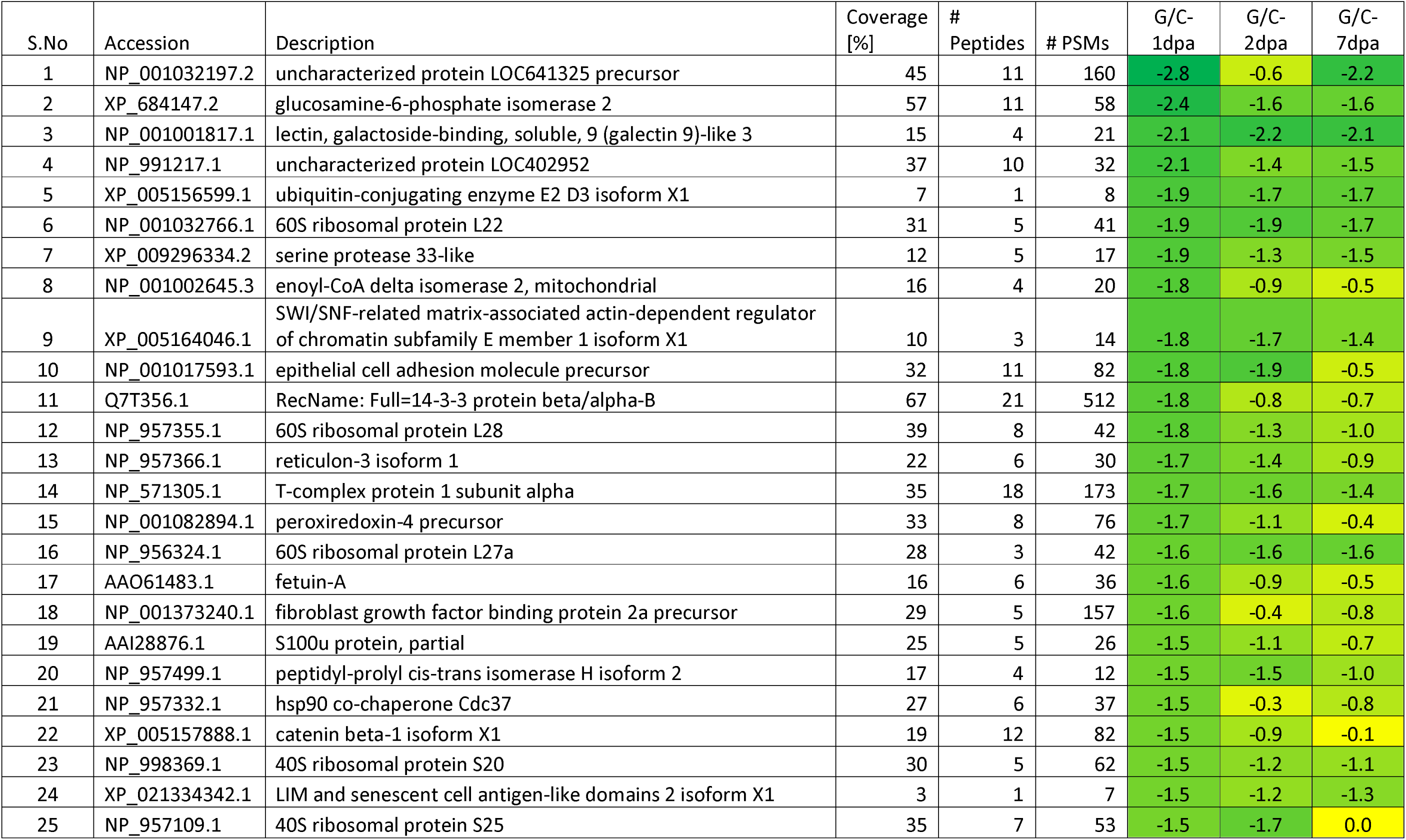
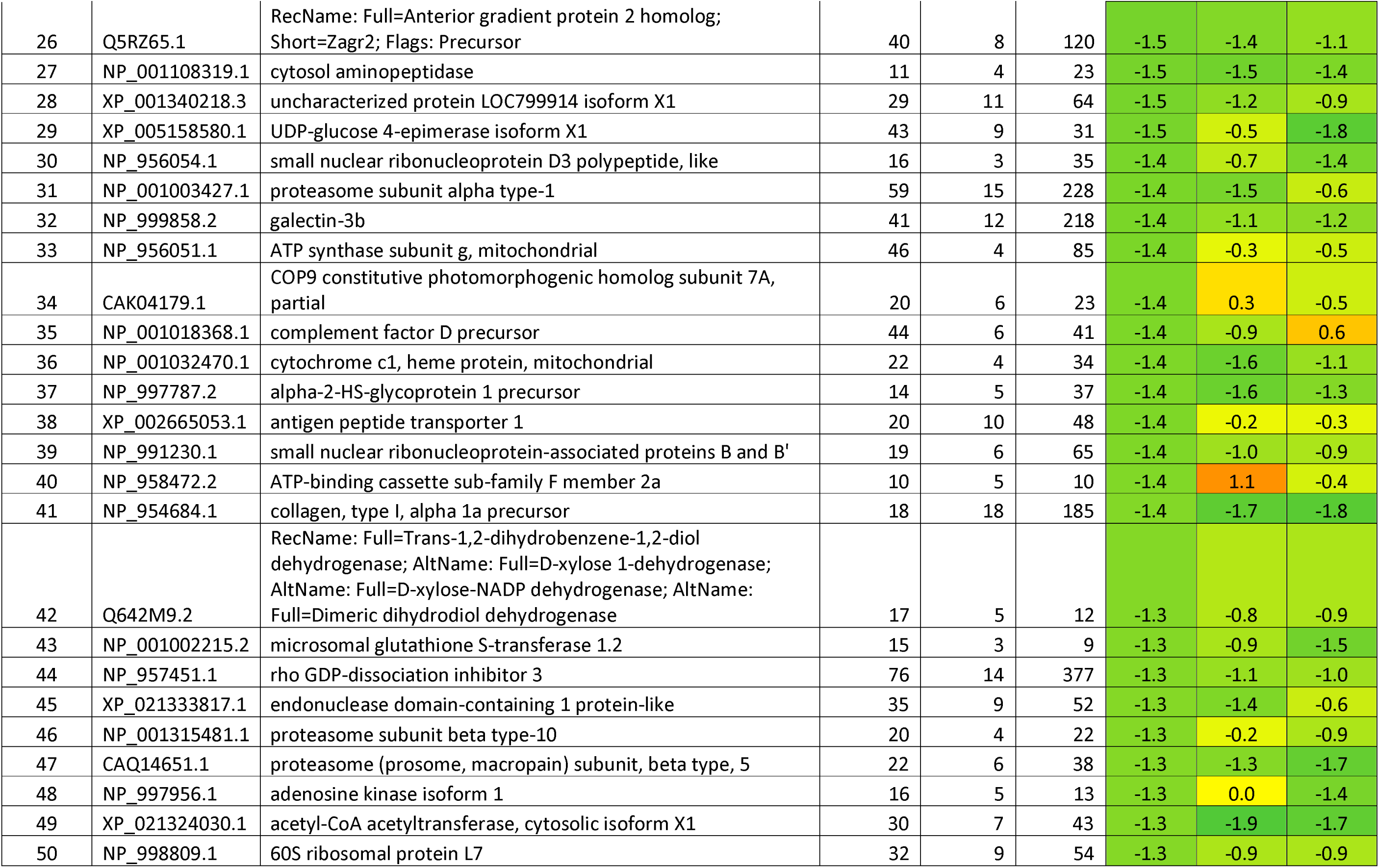
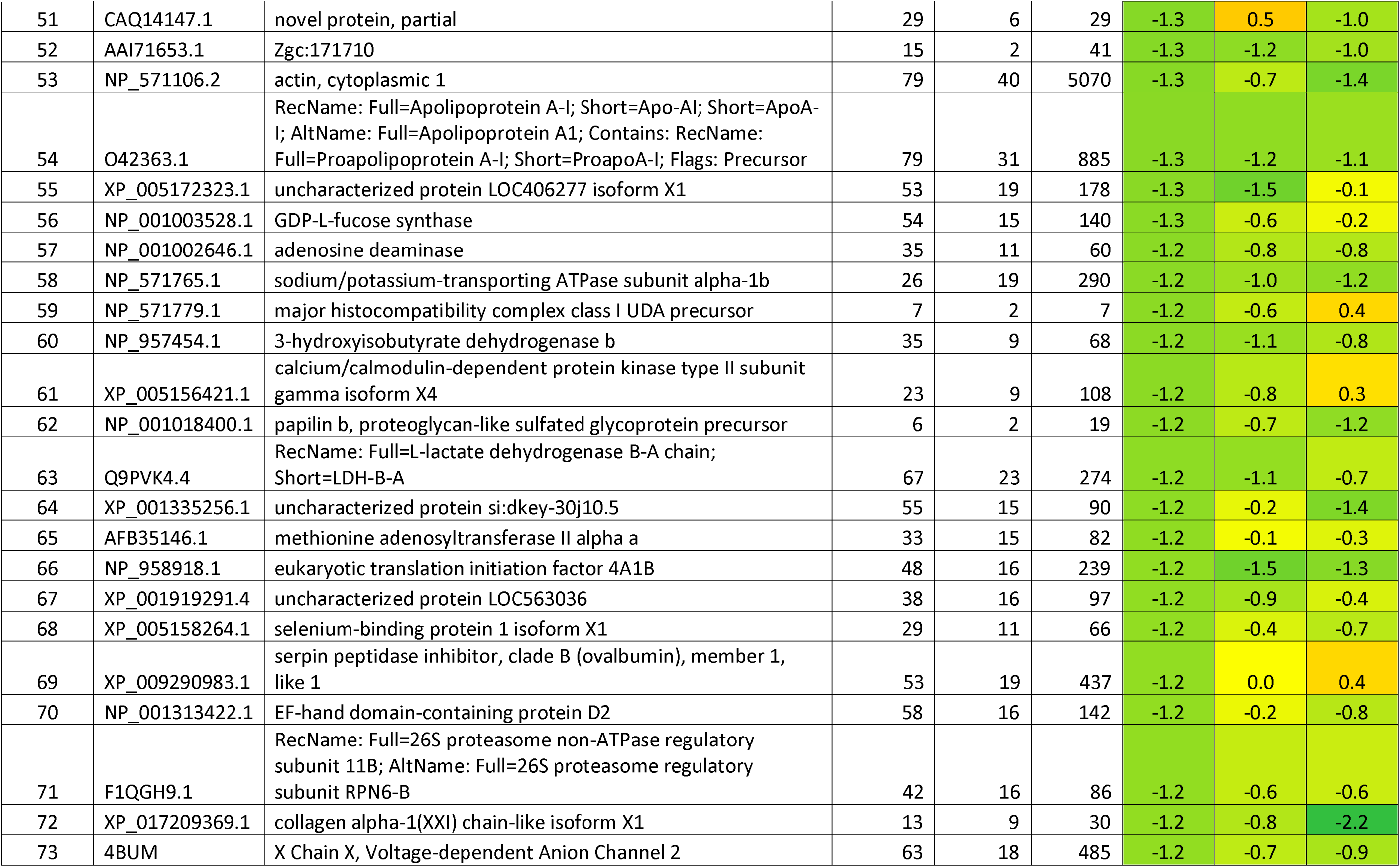
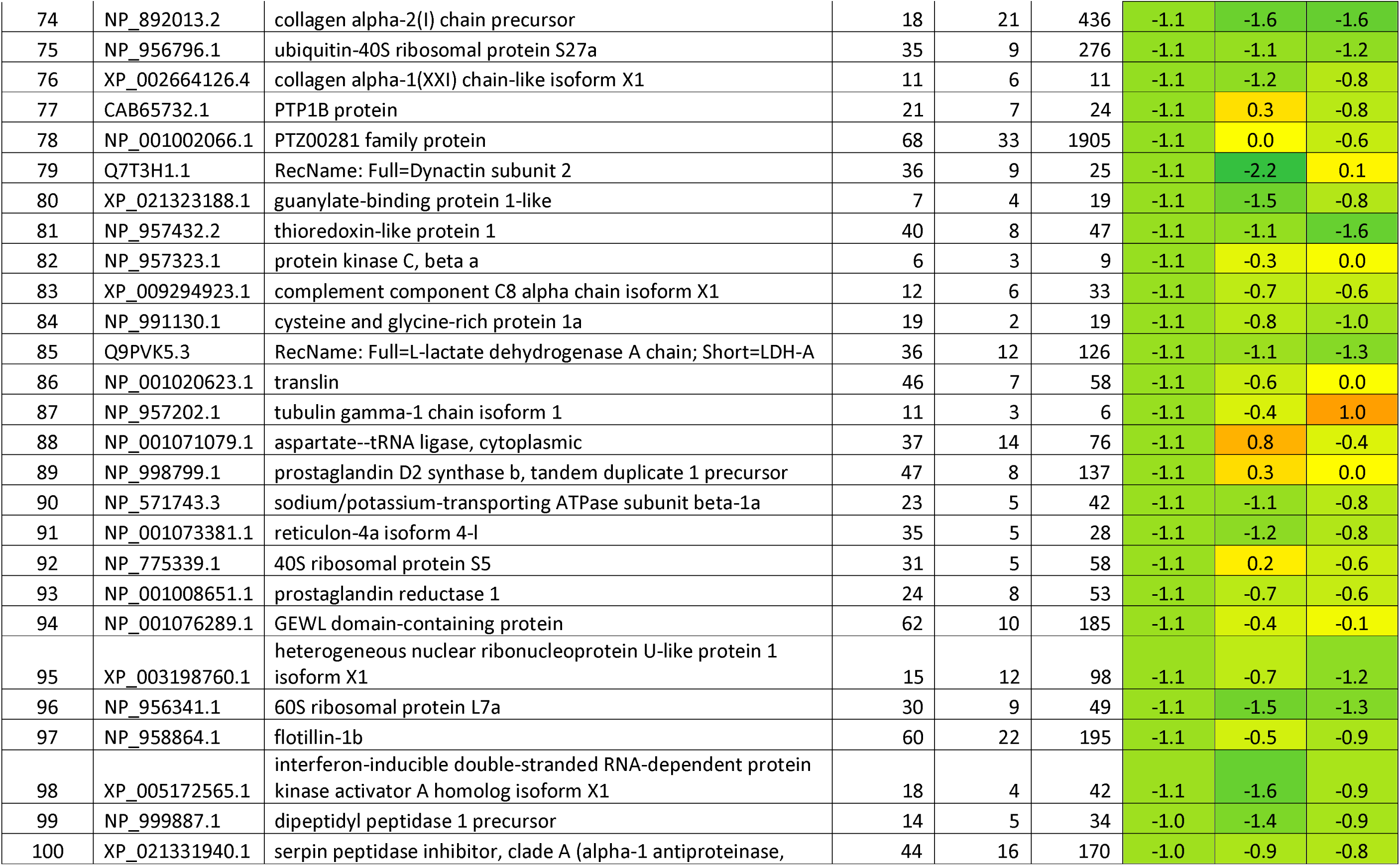
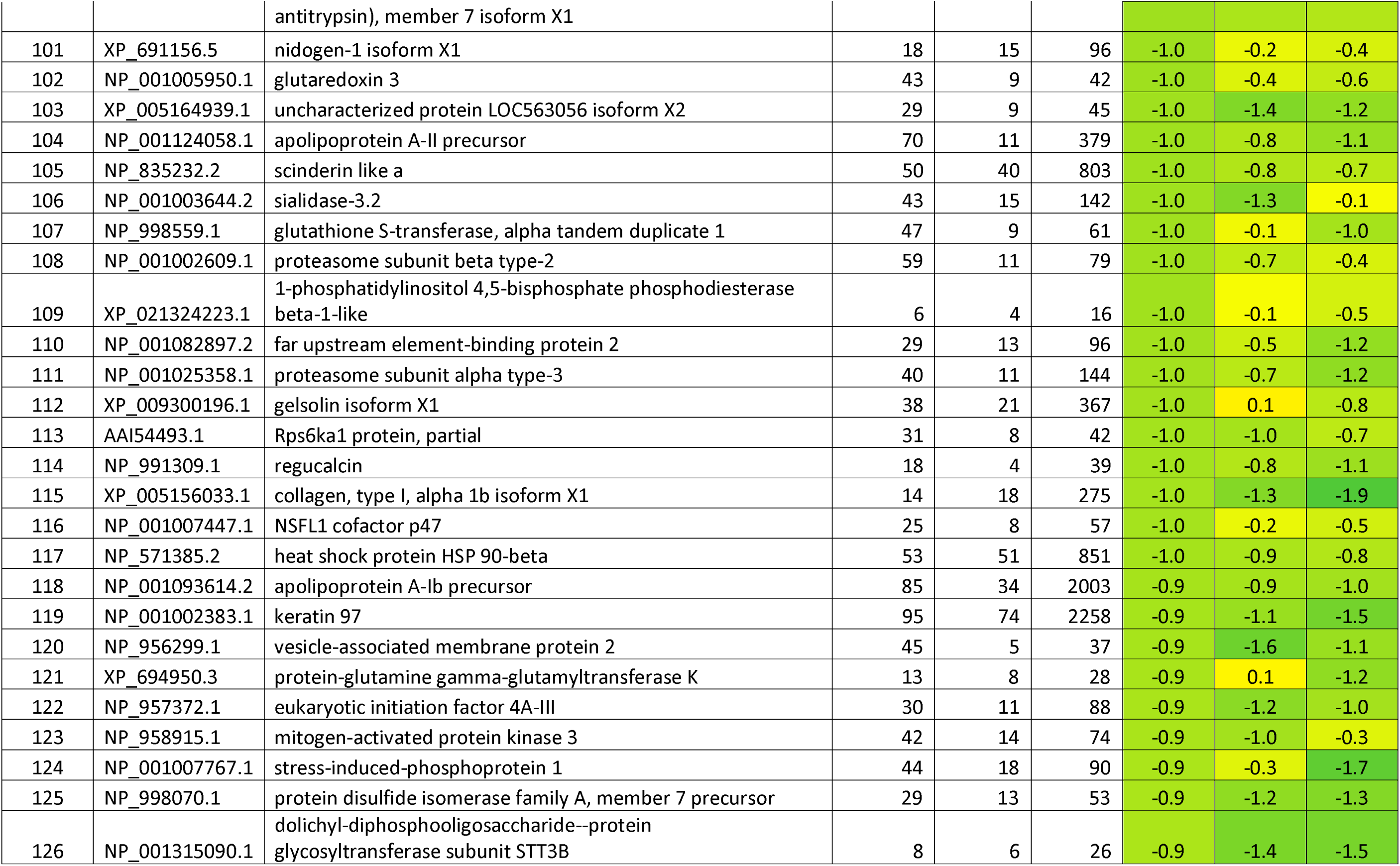
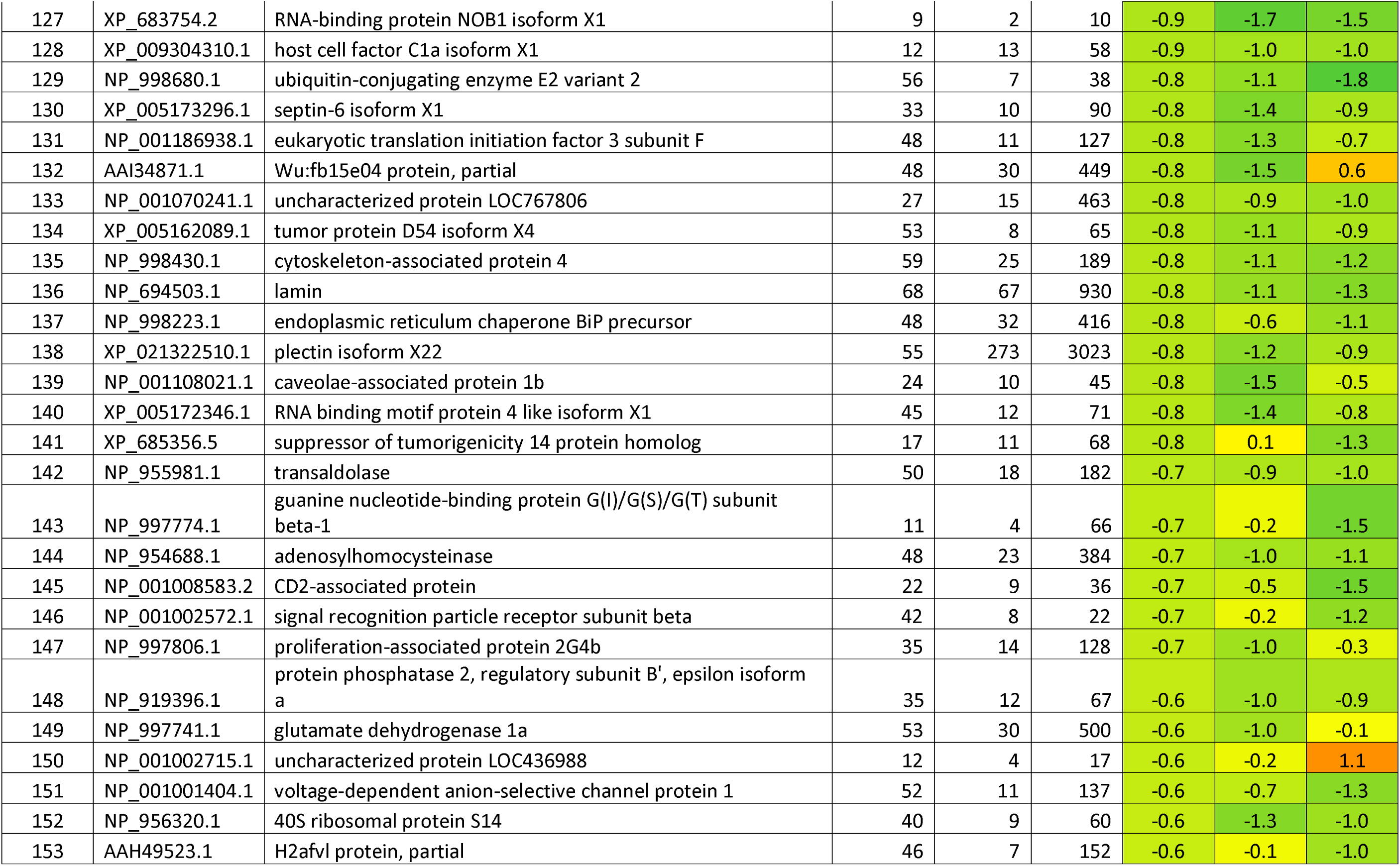
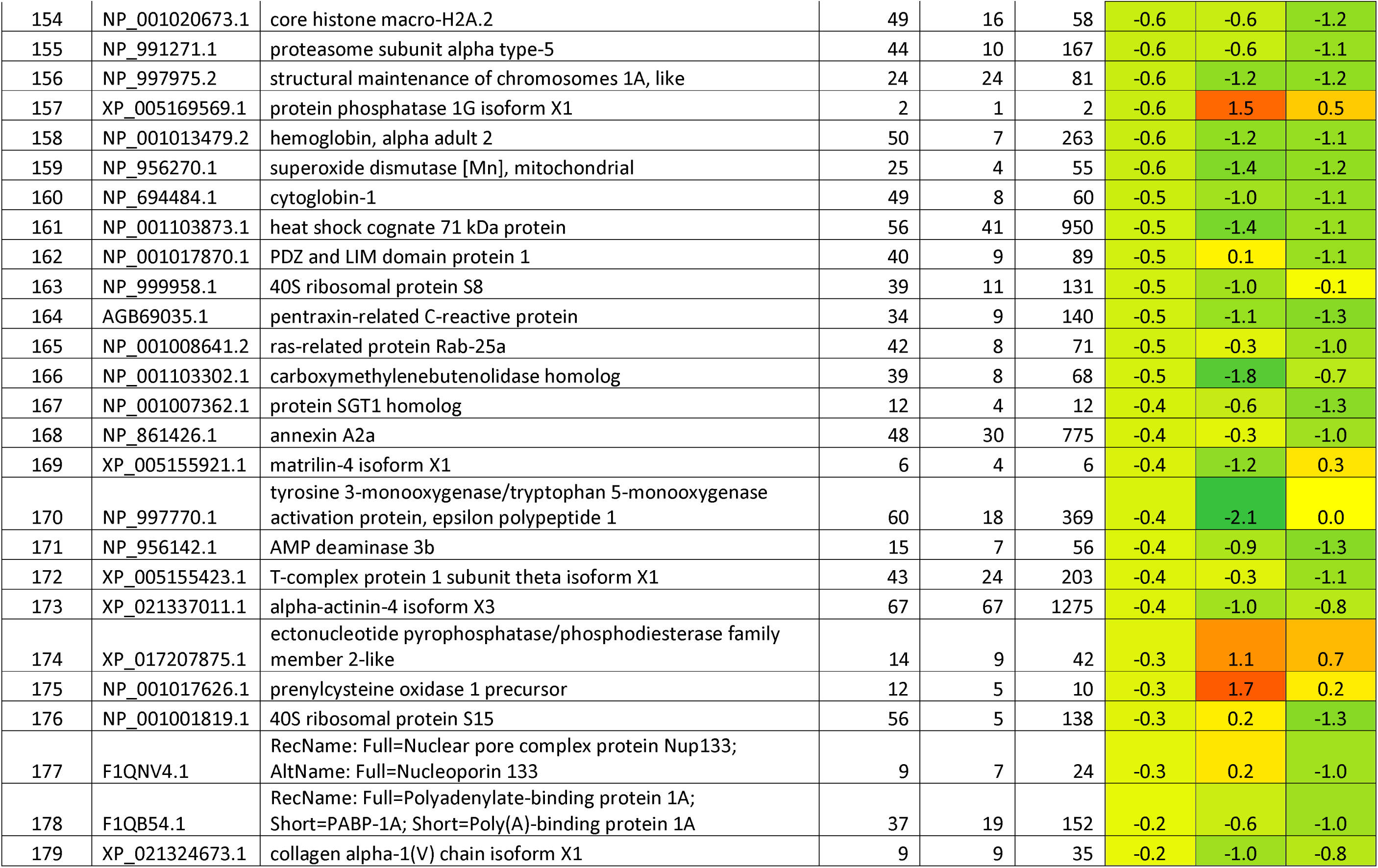
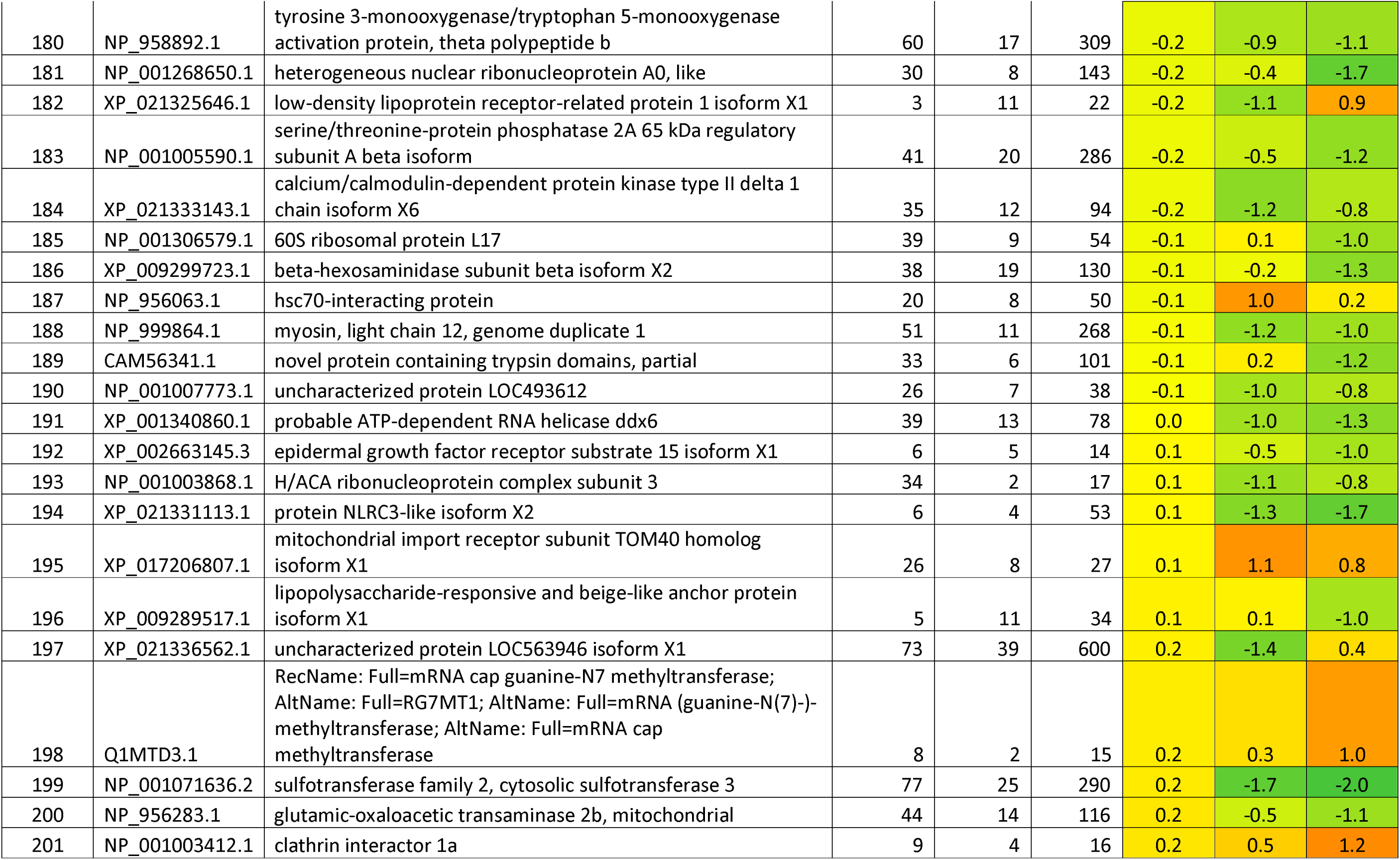
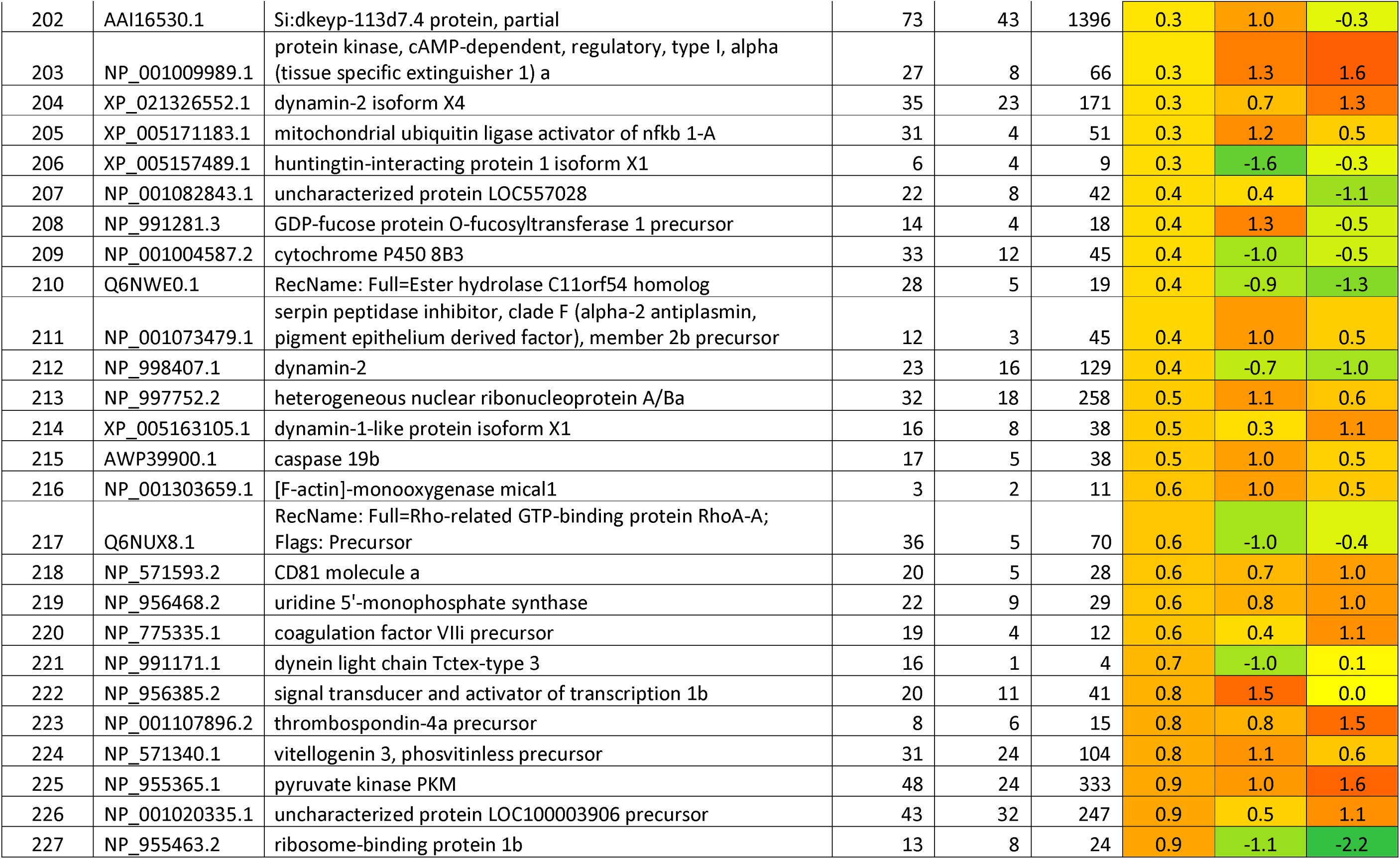
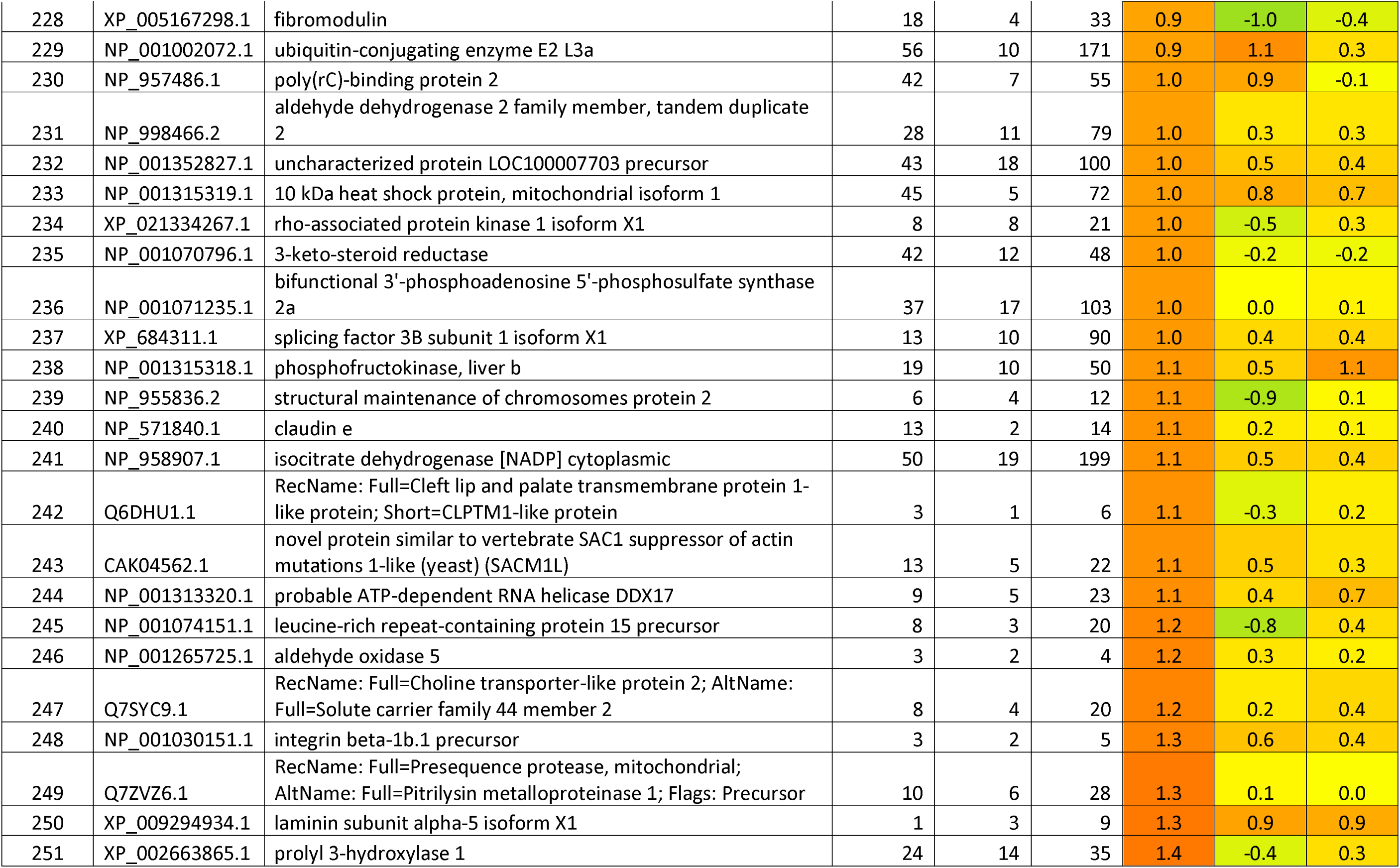
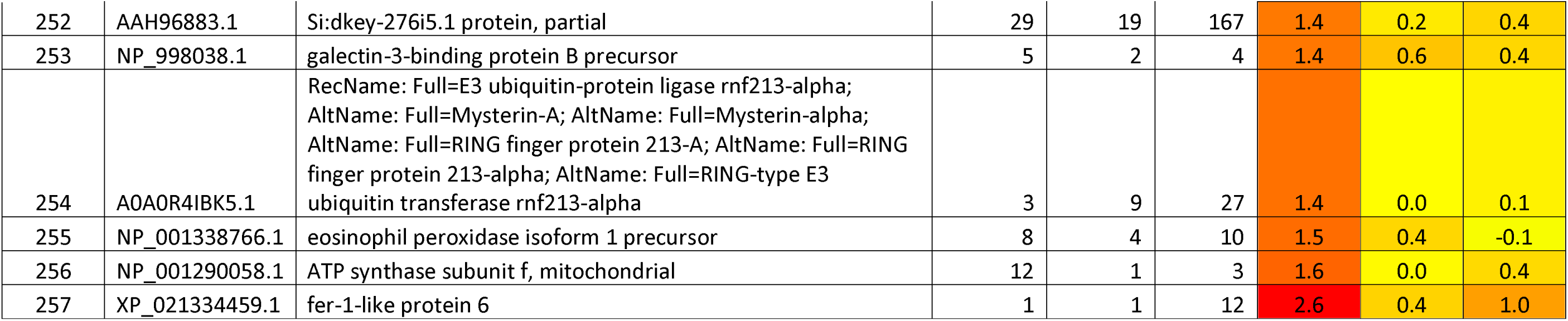
List of Proteins found differentially expressed in zebrafish caudal fin tissue upon exposure to Glucose.

**Table 3:**
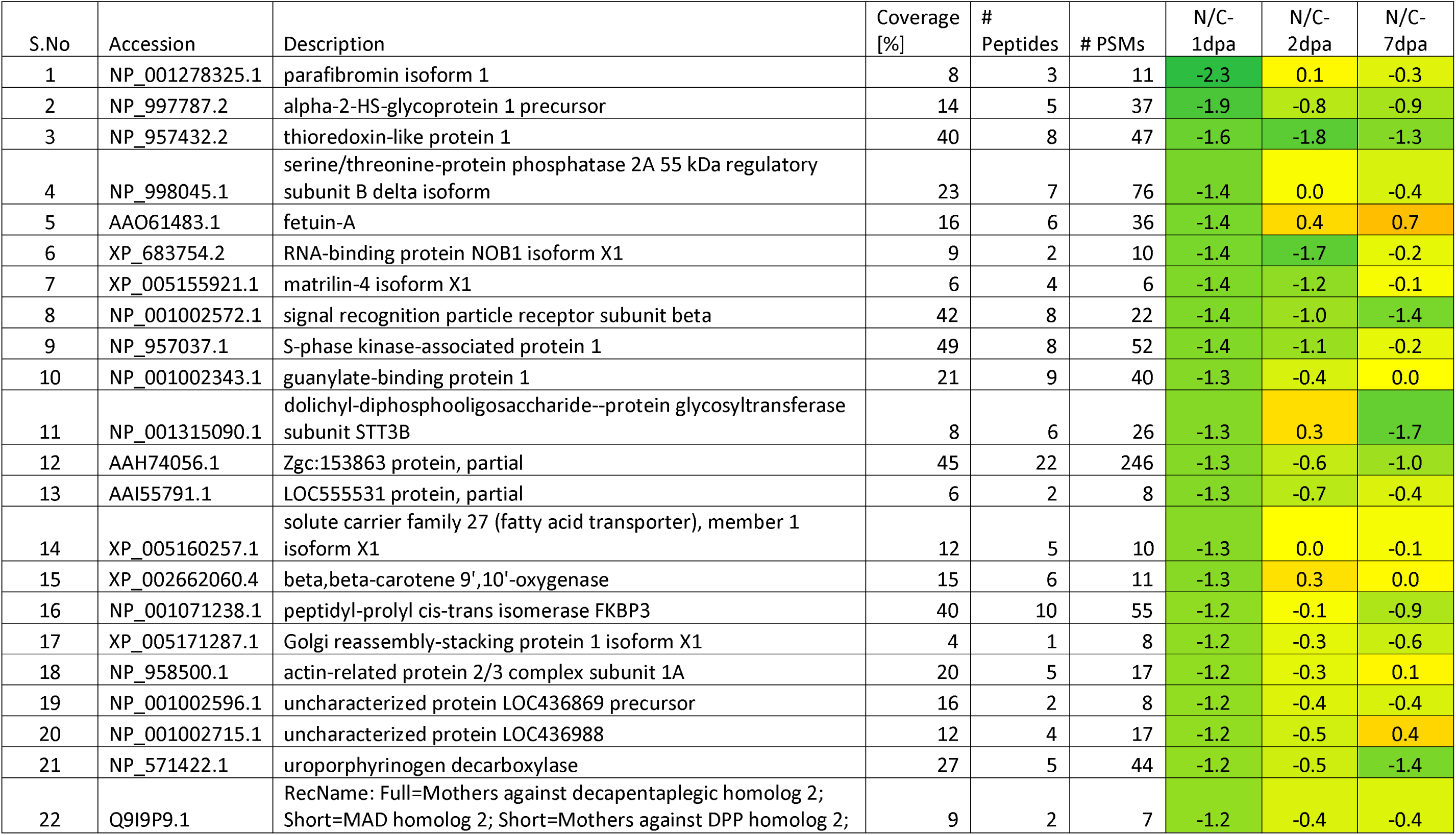
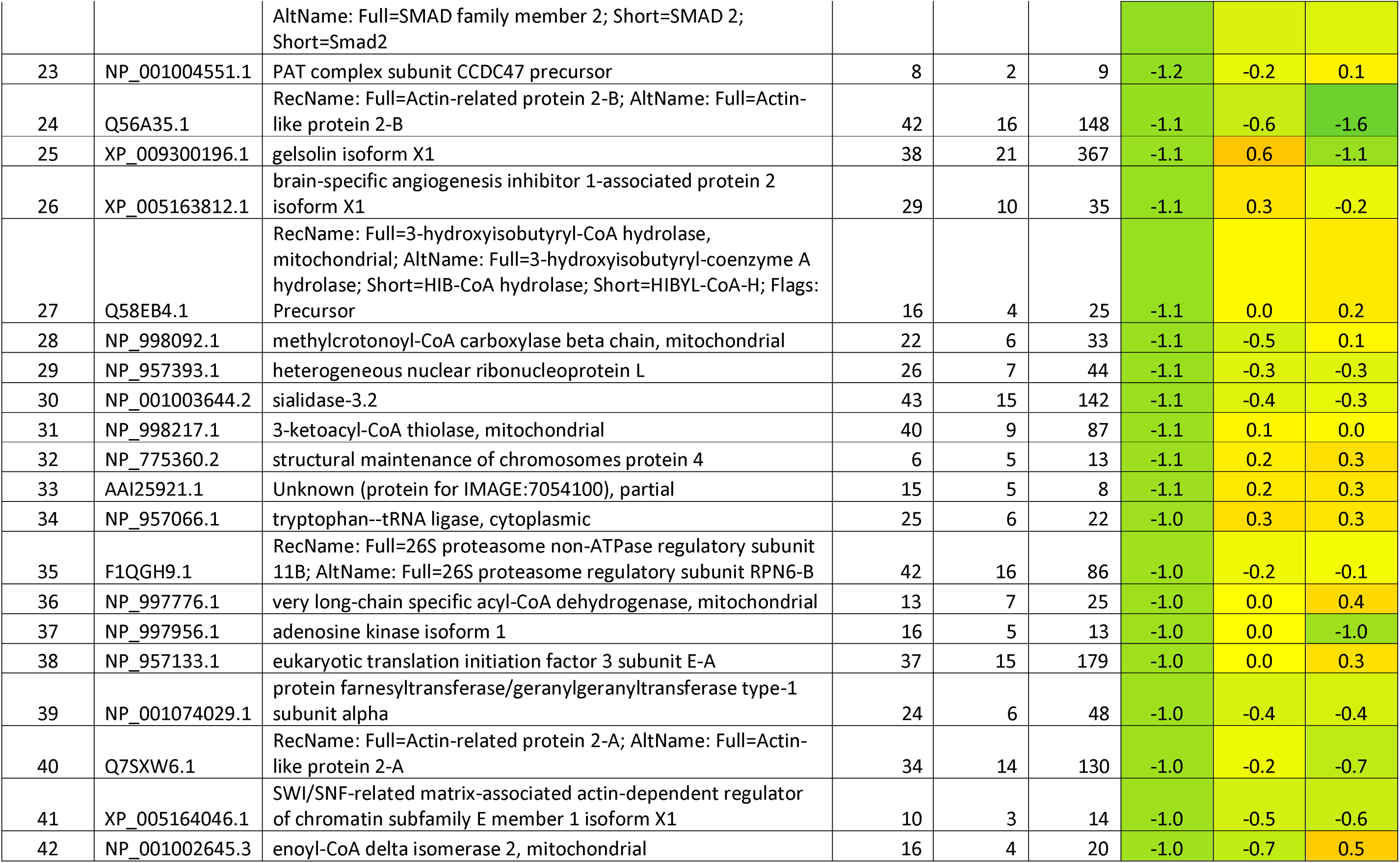
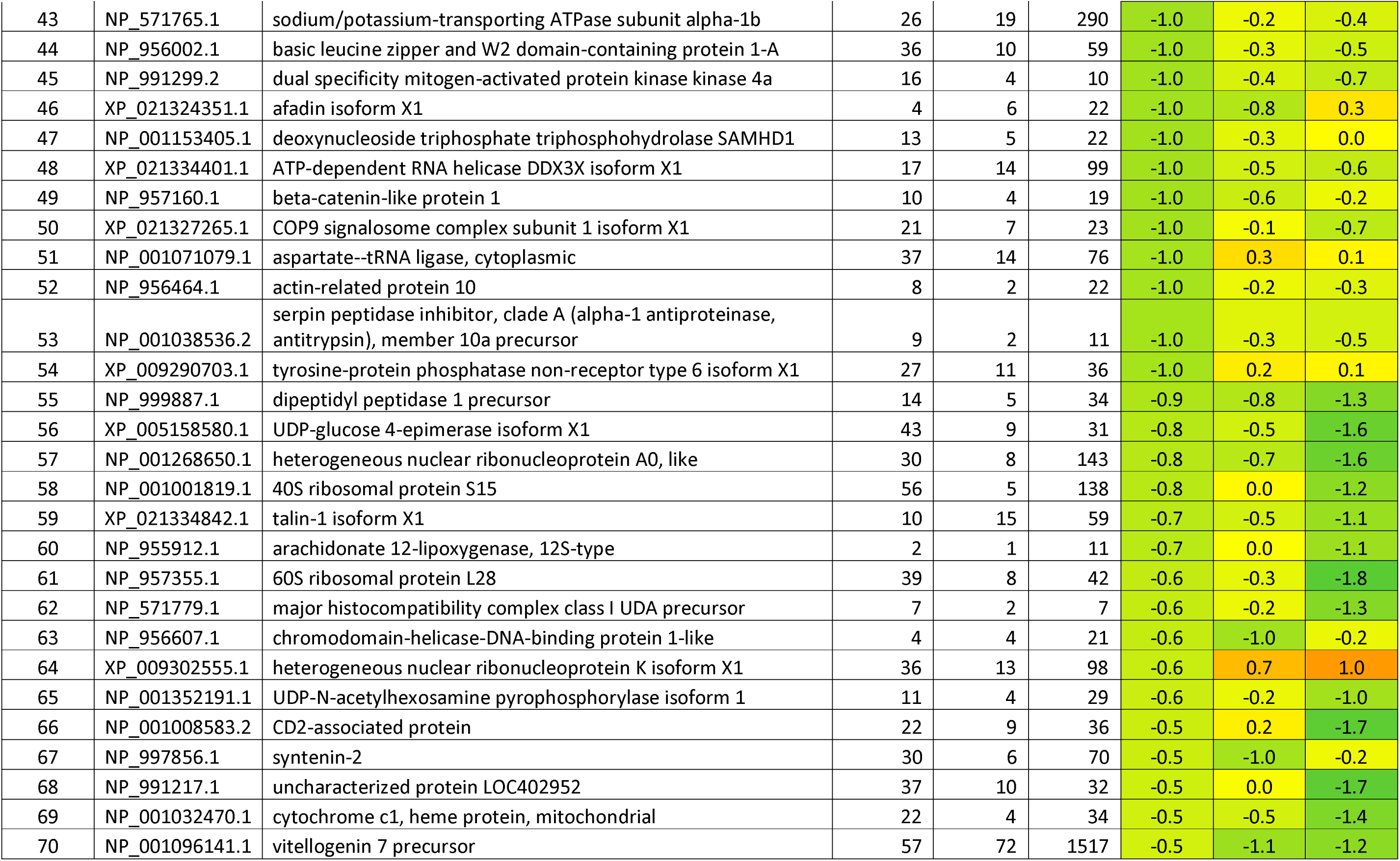
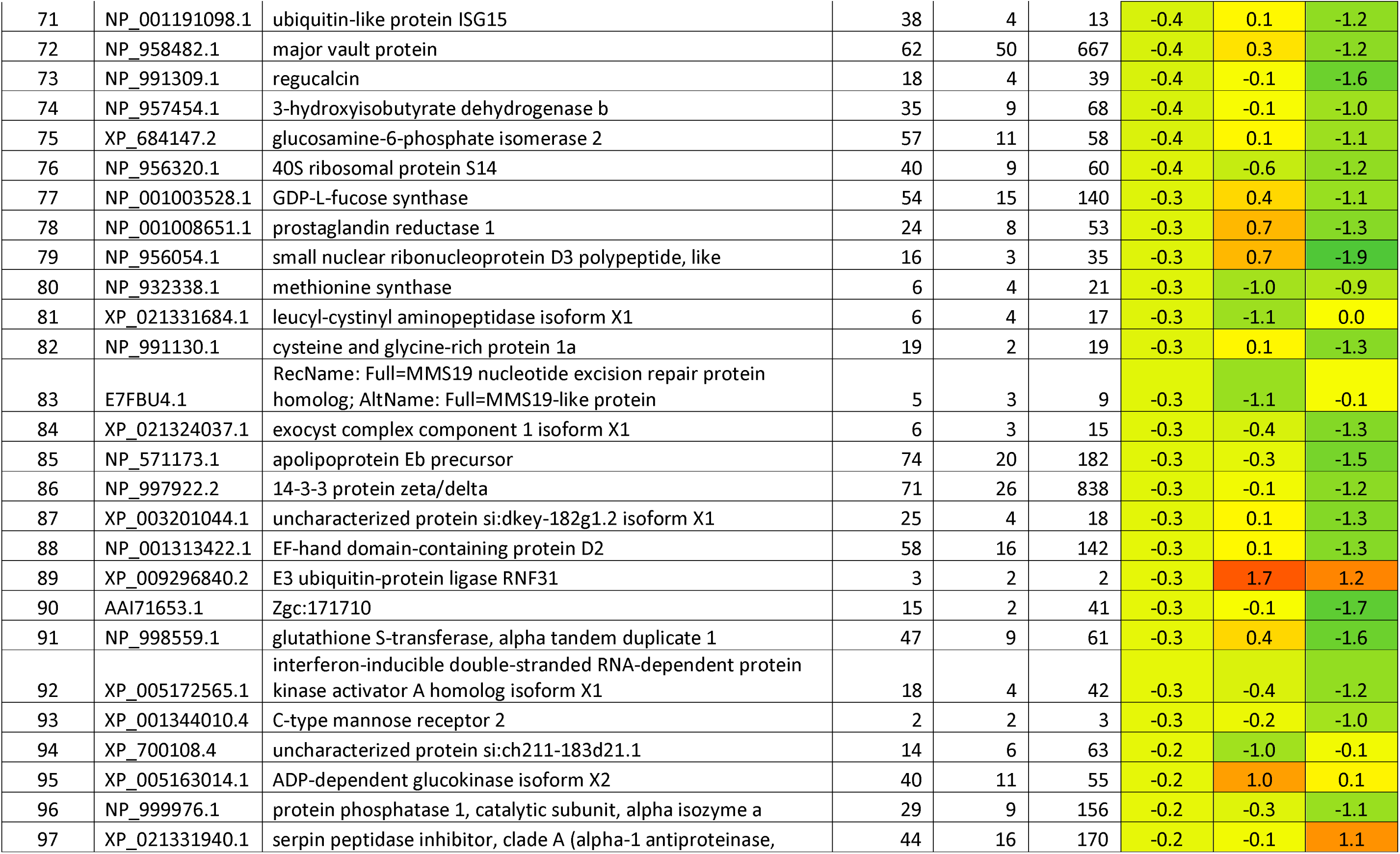
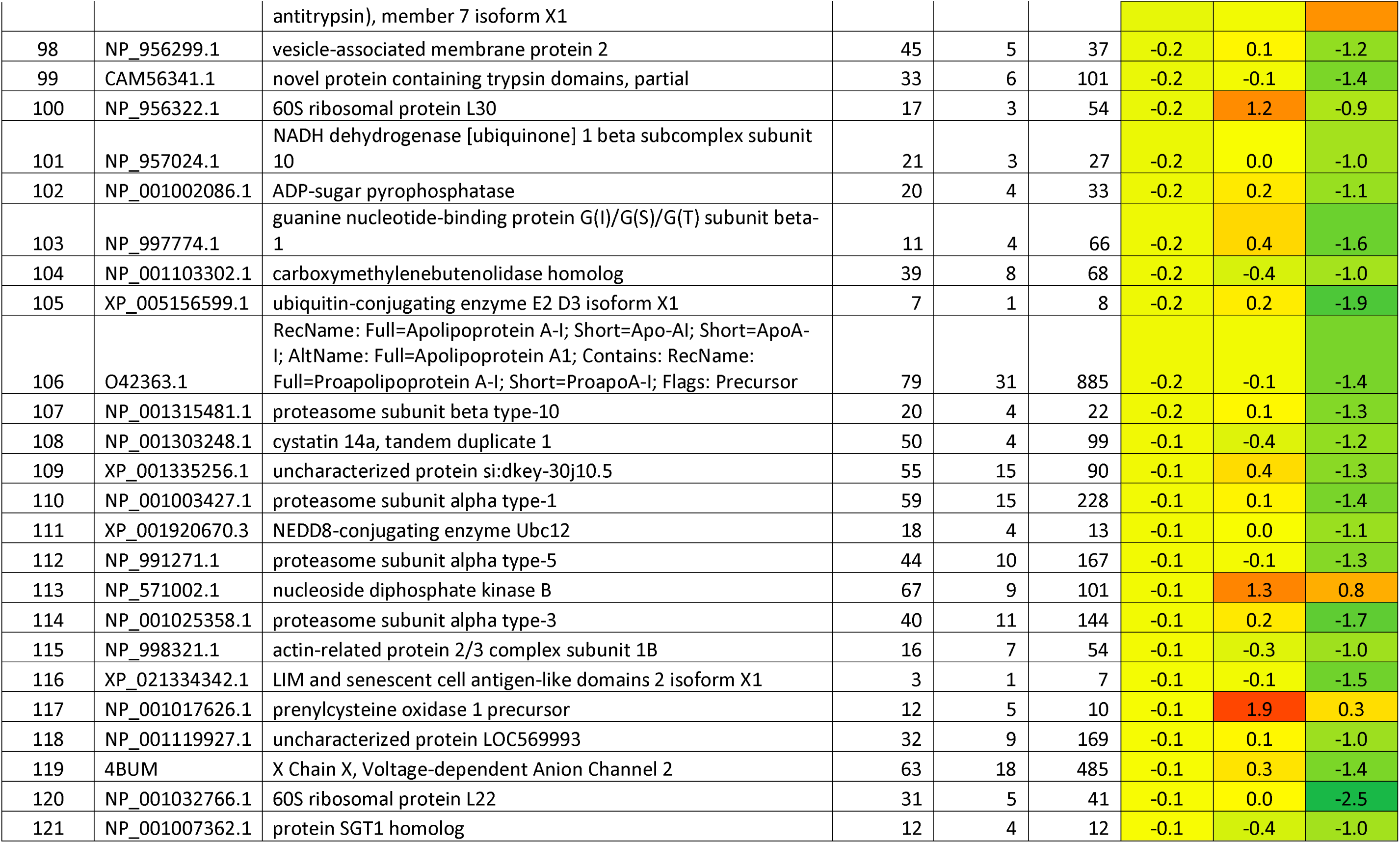
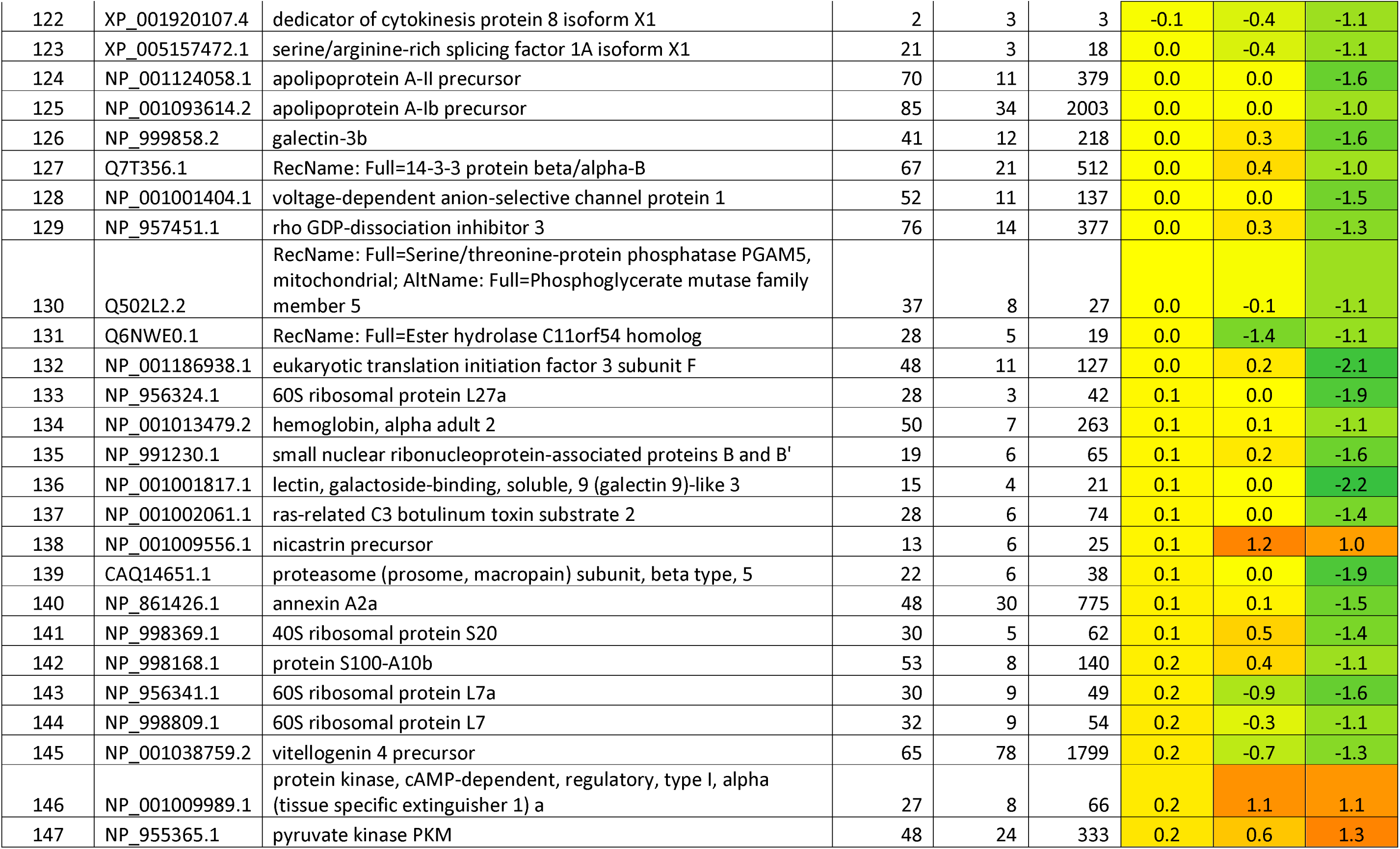
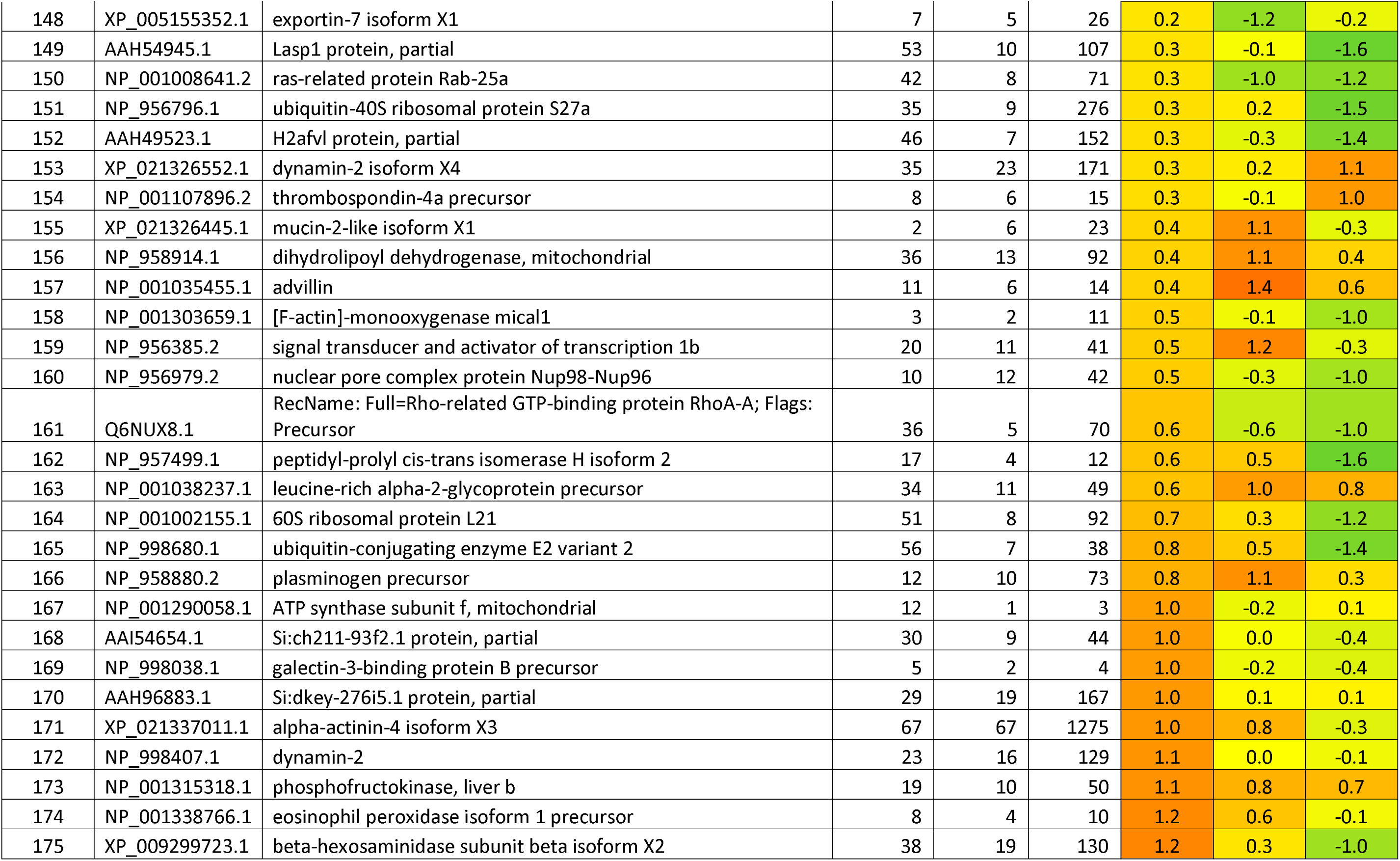
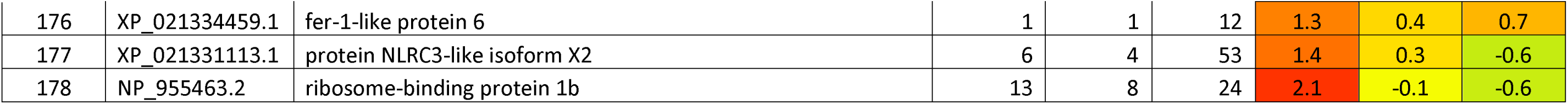
List of Proteins found differentially expressed in zebrafish caudal fin tissue upon exposure to NaCl.

**Table 4:**
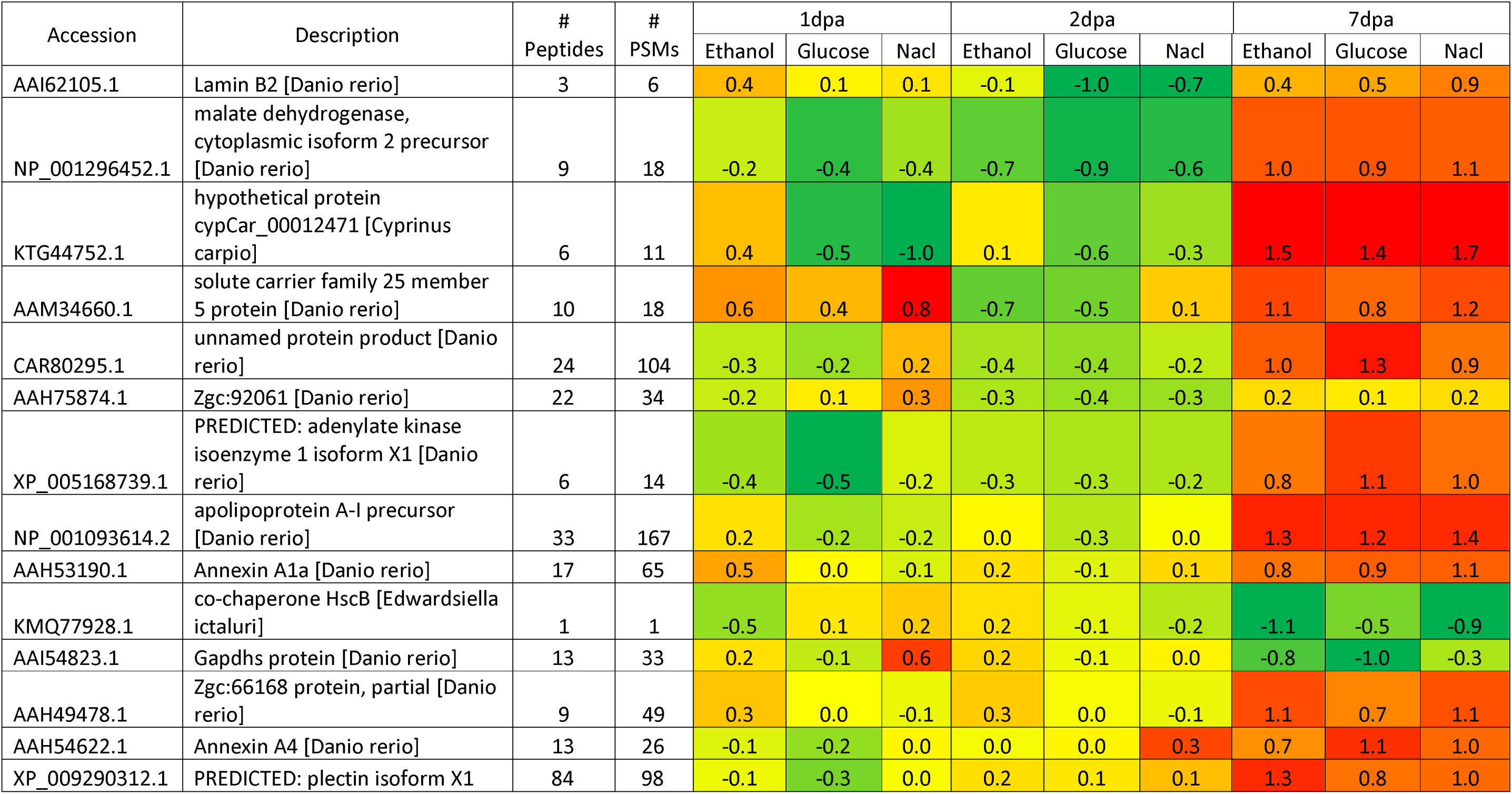
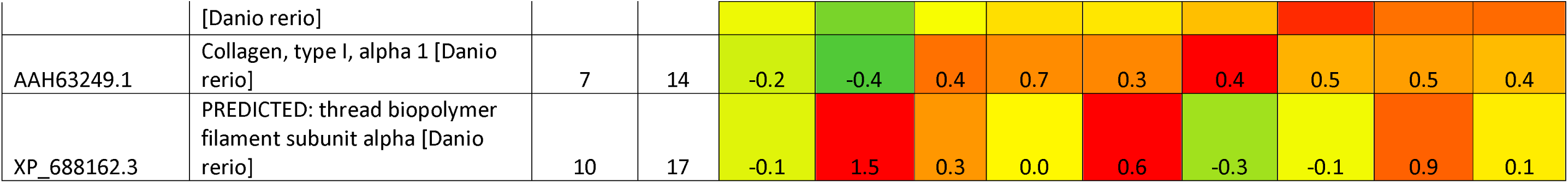
List of Proteins found differentially expressed in zebrafish caudal fin tissue at 1dpa, 2dpa and 7dpa upon exposure to Ethanol, Glucose and NaCl.

| Accession | Description | #<br>Peptides | #<br>PSMs | 1dpa |  |  | 2dpa |  |  | 7dpa |  |  |
| --- | --- | --- | --- | --- | --- | --- | --- | --- | --- | --- | --- | --- |
|  |  |  |  | Ethanol | Glucose | Nacl | Ethanol | Glucose | Nacl | Ethanol | Glucose | Nacl |
| AAI62105.1 | Lamin B2 [Danio rerio] | 3 | 6 | 0.4 | 0.1 | 0.1 | -0.1 | -1.0 | -0.7 | 0.4 | 0.5 | 0.9 |
| NP_001296452.1 | malate dehydrogenase, cytoplasmic isoform 2 precursor [Danio rerio] | 9 | 18 | -0.2 | -0.4 | -0.4 | -0.7 | -0.9 | -0.6 | 1.0 | 0.9 | 1.1 |
| KTG44752.1 | hypothetical protein cypCar_00012471 [Cyprinus carpio] | 6 | 11 | 0.4 | -0.5 | -1.0 | 0.1 | -0.6 | -0.3 | 1.5 | 1.4 | 1.7 |
| AAM34660.1 | solute carrier family 25 member 5 protein [Danio rerio] | 10 | 18 | 0.6 | 0.4 | 0.8 | -0.7 | -0.5 | 0.1 | 1.1 | 0.8 | 1.2 |
| CAR80295.1 | unnamed protein product [Danio rerio] | 24 | 104 | -0.3 | -0.2 | 0.2 | -0.4 | -0.4 | -0.2 | 1.0 | 1.3 | 0.9 |
| AAH75874.1 | Zgc:92061 [Danio rerio] | 22 | 34 | -0.2 | 0.1 | 0.3 | -0.3 | -0.4 | -0.3 | 0.2 | 0.1 | 0.2 |
| XP_005168739.1 | PREDICTED: adenylate kinase isoenzyme 1 isoform X1 [Danio rerio] | 6 | 14 | -0.4 | -0.5 | -0.2 | -0.3 | -0.3 | -0.2 | 0.8 | 1.1 | 1.0 |
| NP_001093614.2 | apolipoprotein A-I precursor [Danio rerio] | 33 | 167 | 0.2 | -0.2 | -0.2 | 0.0 | -0.3 | 0.0 | 1.3 | 1.2 | 1.4 |
| AAH53190.1 | Annexin A1a [Danio rerio] | 17 | 65 | 0.5 | 0.0 | -0.1 | 0.2 | -0.1 | 0.1 | 0.8 | 0.9 | 1.1 |
| KMQ77928.1 | co-chaperone HscB [Edwardsiella ictaluri] | 1 | 1 | -0.5 | 0.1 | 0.2 | 0.2 | -0.1 | -0.2 | -1.1 | -0.5 | -0.9 |
| AAI54823.1 | Gapdhs protein [Danio rerio] | 13 | 33 | 0.2 | -0.1 | 0.6 | 0.2 | -0.1 | 0.0 | -0.8 | -1.0 | -0.3 |
| AAH49478.1 | Zgc:66168 protein, partial [Danio rerio] | 9 | 49 | 0.3 | 0.0 | -0.1 | 0.3 | 0.0 | -0.1 | 1.1 | 0.7 | 1.1 |
| AAH54622.1 | Annexin A4 [Danio rerio] | 13 | 26 | -0.1 | -0.2 | 0.0 | 0.0 | 0.0 | 0.3 | 0.7 | 1.1 | 1.0 |
| XP_009290312.1 | PREDICTED: plectin isoform X1 | 84 | 98 | -0.1 | -0.3 | 0.0 | 0.2 | 0.1 | 0.1 | 1.3 | 0.8 | 1.0 |
|  | [Danio rerio] |  |  |  |  |  |  |  |  |  |  |  |
| AAH63249.1 | Collagen, type I, alpha 1 [Danio rerio] | 7 | 14 | -0.2 | -0.4 | 0.4 | 0.7 | 0.3 | 0.4 | 0.5 | 0.5 | 0.4 |
| XP_688162.3 | PREDICTED: thread biopolymer filament subunit alpha [Danio rerio] | 10 | 17 | -0.1 | 1.5 | 0.3 | 0.0 | 0.6 | -0.3 | -0.1 | 0.9 | 0.1 |

Among the ethanol-treated samples, several proteins were upregulated, including Leucine-rich alpha-2 glycoprotein precursor, pyruvate kinase, vitellogenin 3, neuroblast differentiation-associated protein AHNAK-like isoform X1, F-actin monooxygenase MICAL 1, and plakophilin-3 isoform X1, These findings suggest their roles in cellular metabolism, structural integrity, and regenerative mechanisms. Conversely, proteins such as exportin-5, C-reactive protein 3 precursor, desmoplakin-like, cadherin-like protein 26, heat shock protein HSP 90-beta, major vault protein, and phospholipase C delta 1b isoform X1 were significantly downregulated, indicating potential suppression of certain regulatory pathways during ethanol-induced regeneration impairment (Table1).

In the glucose-exposed samples, proteins such as complement factor D precursor, serpin peptidase inhibitor clade B (ovalbumin), low-density lipoprotein receptor-related protein 1 isoform X1, clathrin interactor 1a, dynamin-2 isoform X4, and mitochondrial ubiquitin ligase activator of NF-kB 1-A exhibited upregulation. These proteins are involved in immune response modulation, lipid metabolism, and cytoskeletal reorganization. Meanwhile, proteins like glucosamine-6-phosphate isomerase 2, 60S ribosomal protein L22, peroxiredoxin-4 precursor, and catenin beta-1 isoform X1 were downregulated, reflecting possible disruptions in protein synthesis and oxidative stress responses (Table 2).

NaCl exposure led to the upregulation of proteins such asfetuin-A, actin-related protein 2/3 complex subunit 1A, PAT complex subunit CCDC47 precursor, structural maintenance of chromosomes protein 4, tryptophan--tRNA ligase, and enoyl-CoA delta isomerase 2. Proteins that were downregulated included dipeptidyl peptidase 1 precursor, UDP-glucose 4-epimerase isoform X1, heterogeneous nuclear ribonucleoprotein A0-like, 40S ribosomal protein S15, major histocompatibility complex class I UDA precursor, vitellogenin 7 precursor, and apolipoproteinEb precursor (Table 3).

To validate the findings from the label-free proteomics, we employed the iTRAQ-based quantitative proteomics approach, which confirmed the differential expression of 16 common proteins across all three experimental groups (Table 4). Notably, proteins such as Lamin B2, malate dehydrogenase (cytoplasmic isoform 2 precursor), solute carrier family 25 member 5 protein, Annexin A1a, Annexin A4, and Collagen type I alpha 1exhibited a gradual upregulation as regeneration progressed, These findings hint at their involvement in tissue repair and structural remodelling. In contrast, co-chaperone HscB and Gapdhs protein showed a progressive downregulation, potentially indicating that they do play a role in early stress responses which becomes reduced in later regenerative stages.

These proteins may represent core molecular players involved in the shared regenerative response or stress-adaptive mechanisms triggered by ethanol, glucose, and NaCl treatments. Their differential expression underscores the complexity of the regenerative process and highlights key candidates for further functional validation in future studies. Taken together, our proteomic analyses demonstrate that exposure to different small molecules induces unique and overlapping protein expression changes, with potential implications for regenerative mechanisms and cellular stress responses in zebrafish caudal fin tissue.

A total of 327 genes/proteins were found mapped by the Ingenuity Pathway Analysis (IPA) software for various canonical pathways as well as disease and function networks concerning each of the treated groups. Several cell-related diseases and functions, majorly of cell movement, cell viability, cell survival, migration of cells apart from carcinoma and various tumors were identified based on the differential protein expression of the treated groups. The most significant canonical pathways that were found to be associated with differentially regulated proteins from proteomics analysis were the GP6 Signaling pathway, Mitochondrial dysfunction, RHO GTPase cycle, Class I MHC mediated antigen signaling, S Phase, Deubiquitination, TCR signaling and various other metabolic pathways.

The IPA analysis of disease and functions in Ethanol, Glucose, and NaCl-treated groups revealed distinct regulation patterns (figure 3). Across all groups, organismal death was the only biological activity consistently upregulated. In the ethanol group, it was initially downregulated but showed significant upregulation by 7 days post-amputation (dpa). In contrast, both NaCl and glucose treatments showed persistent upregulation at all time points. For the ethanol-treated group, biological activities such as cell engulfment, cell movement, migration, viability, survival, endocytosis, phagocytosis, and various cancer-related activities, including metastasis, extracranial solid tumor, lung cancer and invasive cancer, were initially upregulated but showed substantial downregulation by 7dpa.

**Figure 3:**
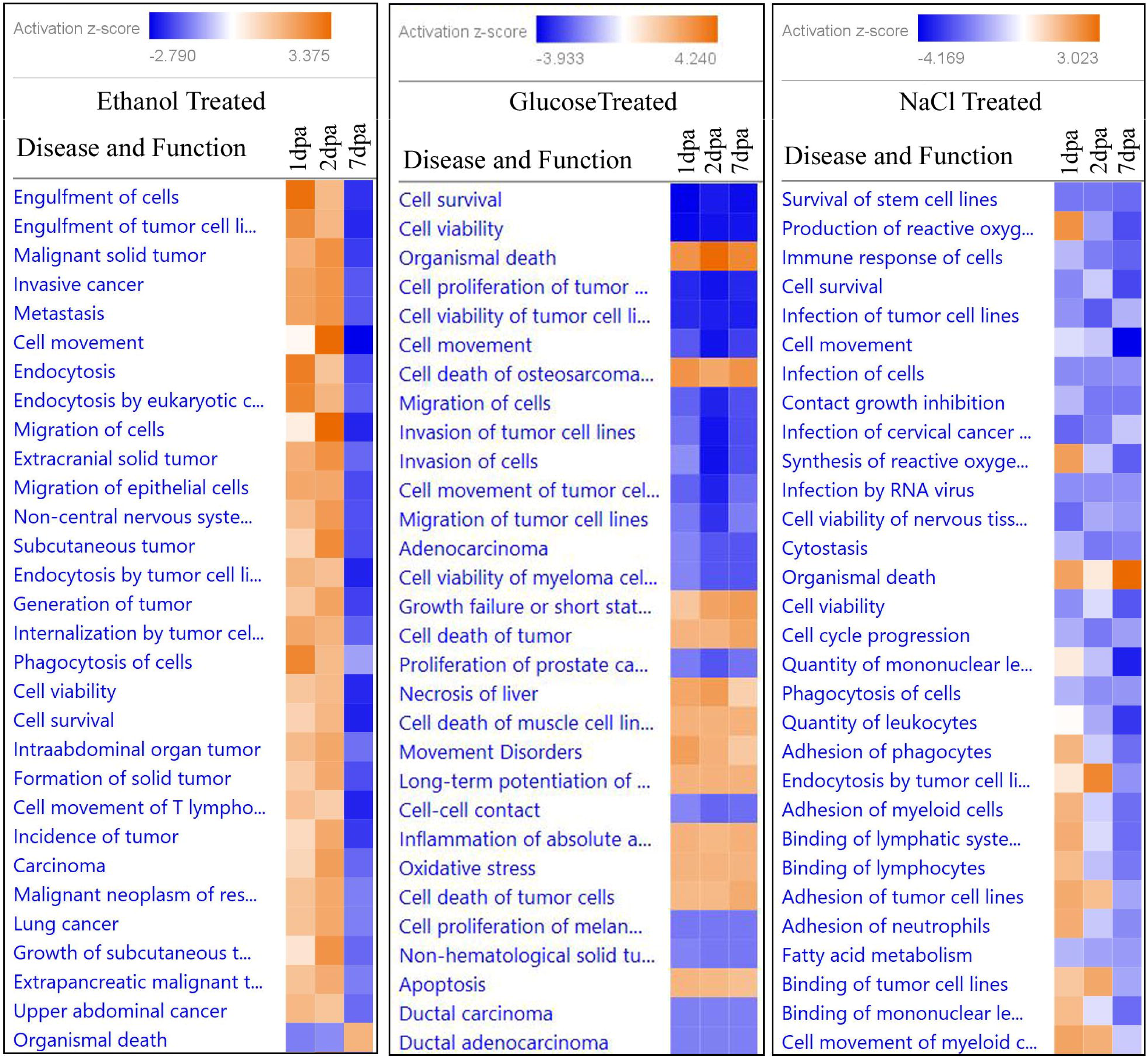
Disease & Functions associated with differentially expressed proteins based on IPA analysis for zebrafish caudal fin regeneration under ethanol, glucose, and NaCl treatment.

The glucose-treated group showed distinct trends, with activities such as osteosarcoma cell death, muscle cell line death, tumor growth failure, liver necrosis, movement disorders, oxidative stress, and apoptosis being upregulated consistently across time points. Conversely, activities related to ductal carcinoma, adenocarcinoma, prostate cancer proliferation, tumor invasion and migration, and cell viability were downregulated at all time points. In the NaCl-treated group, activities related to fatty acid metabolism, leukocyte quantity, RNA virus infection, tumor cell infection, stem cell survival, immune response, cell viability, cytostasis, and cell cycle progression were consistently downregulated across all time points. However, activities linked to reactive oxygen synthesis, cell adhesion (involving phagocytes, neutrophils, and myeloid cells), and binding interactions (e.g., lymphocyte binding and tumor cell line interactions) showed initial upregulation but diminished by 7dpa. These patterns highlight how each treatment influences specific biological activities in tissue repair and injury response mechanisms.

The IPA analysis of canonical pathways across the Ethanol, Glucose, and NaCl-treated groups showed distinct pathway activation patterns (figure 4). For the glucose-treated group, all identified pathways, including S phase, Deubiquitination, Neddylation, RAF/MAP kinase cascade, TCF-dependent signaling, Hedgehog ligand biogenesis, Hedgehog on state, and Signaling by ROBO receptors exhibited a consistent downregulation across all time points. This suggests an inhibition of cell cycle progression, signaling, and regulatory mechanisms throughout the treatment. In the ethanol-treated group, a marked upregulation was observed in the Protein sorting and Neutrophil extracellular trap signaling pathways, indicative of enhanced cellular sorting and immune responses early in the regenerative process. Other pathways, including GP6 signaling, Wound healing signaling, Actin-cytoskeleton signaling, Hepatic fibrosis signaling, IL-12 signaling, and RHO GTPase cycle, showed a pattern of gradual downregulation by 7 days post-amputation (dpa), indicating a progressive suppression of immune, cytoskeletal, and regenerative responses over time. The NaCl-treated group displayed unique pathway regulation, with ROBO Slit signaling as the only upregulated pathway, suggesting involvement in guidance and migration processes. Other pathways, including Neddylation, Protein ubiquitination, TCF-dependent signaling, RHO GTPase cycle, RAF/MAP kinase cascade, RAC signaling, Deubiquitination, IL-1 family signaling, Integrin signaling, S-phase, KEAP1-NFE2L2 pathway, TCR signaling, and Ephrin receptor signaling, demonstrated downregulation, pointing to diminished cell signaling, immune modulation, and antioxidant responses across time points. These distinct patterns across treatment groups indicate differential regulation of pathways related to cellular sorting, immune response, cytoskeletal dynamics, and signaling, reflecting each treatment’s unique impact on cellular mechanisms and regenerative capacity.

**Figure 4:**
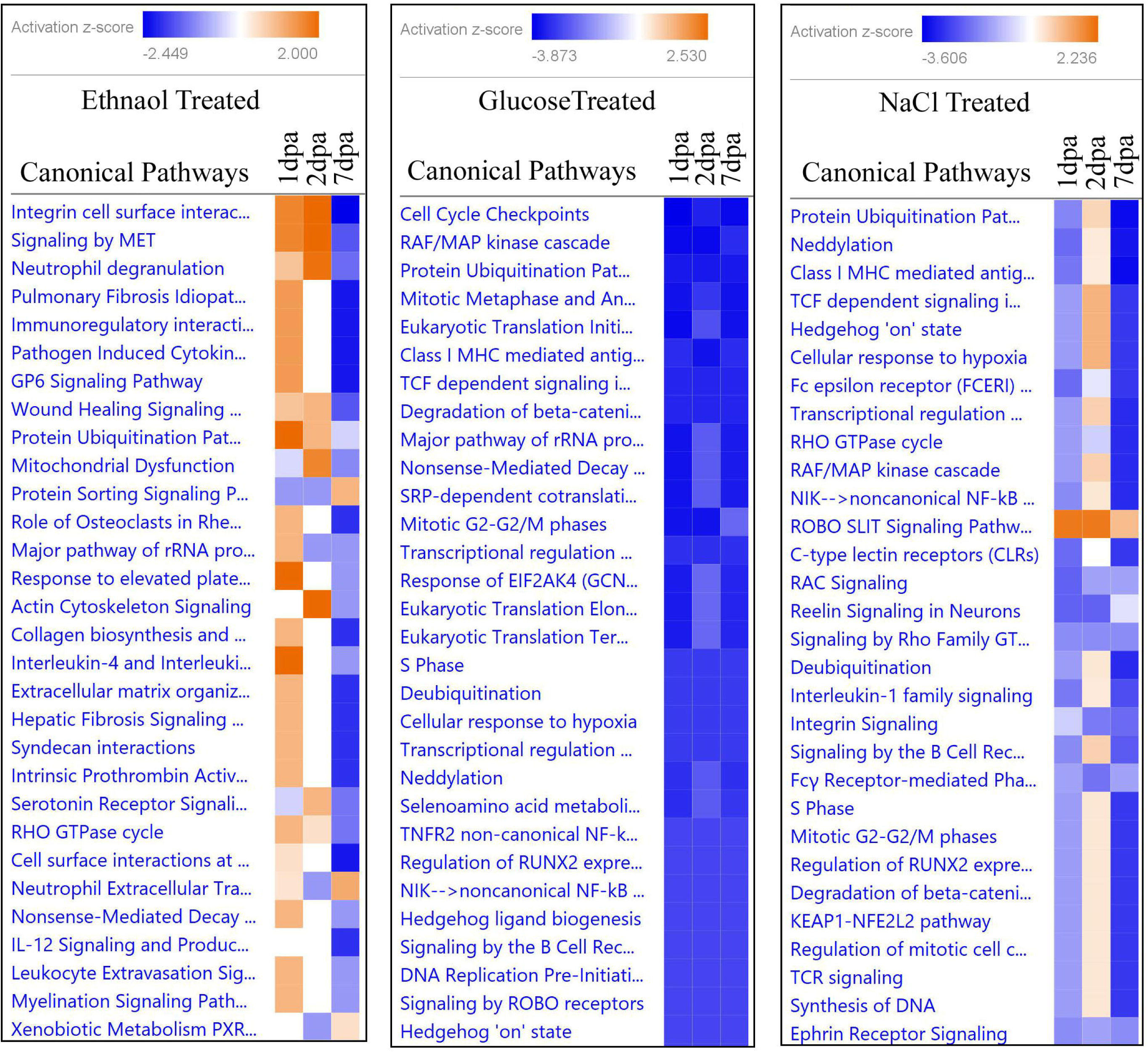
Canonical pathways associated with differentially expressed proteins based on IPA analysis for zebrafish caudal fin regeneration under ethanol, glucose, and NaCl treatment.

## Discussion

In this study, we investigated the effects of small molecules—0.5% ethanol, 1% glucose, and 0.2% NaCl on the regeneration of the adult zebrafish caudal fin as well as in their behaviour. Our findings provide insights into how each molecule influences the regeneration process, potentially through modulation of cellular and molecular pathways essential for wound healing, tissue growth, and differentiation. Our findings indicate that zebrafish can tolerate these concentrations without lethal effects, supporting previous data on their resilience to various chemical exposures at low concentrations (20, 21). Most notable is how each molecule influenced fin regeneration, behavioural stress responses, and proteomic profiles in distinct ways, which suggests unique underlying molecular and cellular mechanisms.

### Impact of Small Molecules on Caudal Fin Regeneration

Upon exposure to 0.5% ethanol, 1% glucose, and 0.2% NaCl, zebrafish showed varied regeneration rates across fin lobes and clefts, with ethanol exerting the most significant inhibitory effect. Ethanol’s effects are complex; at lower concentrations, it can act as a cellular signaling molecule, but at higher levels, it may induce cytotoxicity, disrupt cell signaling, and impact gene expression related to regeneration. Ethanol’s impact on regenerative processes may stem from its ability to alter cellular signaling and stress response pathways, as shown in prior studies that link ethanol exposure to oxidative stress and apoptosis, particularly in developing tissues [20]. Ethanol’s interference with these processes may explain the inhibited regeneration observed in our study, consistent with findings that ethanol can reduce progenitor cell proliferation and differentiation in zebrafish models [22].

Similarly, the glucose-treated group displayed regeneration rates that varied in comparison to the control, underscoring glucose’s complex role in regeneration. Glucose plays a crucial role in cellular energy metabolism, and studies have shown that elevated glucose levels can enhance tissue repair by providing energy for cell proliferation and extracellular matrix formation. Glucose provides essential energy for cellular processes; however, its overabundance can result in osmotic and metabolic stress, potentially hindering regeneration if not balanced correctly [21]. Our findings are consistent with studies suggesting that glucose concentration must be carefully balanced to avoid detrimental effects on cellular processes during regeneration.

Sodium ions play an essential role in cell signaling and osmoregulation, which are critical during the wound healing and regenerative phases. The NaCl group also showed some inhibition in regrowth, suggesting that ionic imbalance impacts cellular homeostasis and may hinder wound healing. However, high NaCl concentrations can lead to osmotic stress, potentially disrupting cellular signaling and osmoregulatory processes, thereby altering cellular activities essential for regeneration [23]. Our findings indicate that NaCl exposure inhibits regenerative outcomes, suggesting that ionic balance is crucial for optimal regeneration. This is in line with previous studies that highlight how ionic composition influences cellular responses during tissue repair, especially in aquatic models like zebrafish.

### Behavioural Analysis and Stress Responses

Our behavioural analysis revealed stress-induced changes, including decreased velocity and distance moved across all treated groups. Of the groups, those that were ethanol-treated displayed the most significant reduction. Fish in treated groups showed a tendency to remain in the lower tank levels, a known stress indicator in zebrafish behaviour [24]. Behavioural stress in zebrafish can result from ethanol exposure, which has been shown to influence neurobehavioral responses by disrupting neural signaling pathways [23]. Our findings are consistent with prior studies demonstrating that zebrafish under stress exhibit lower exploratory behaviour and altered movement patterns [25]. The glucose and NaCl-treated groups showed similar stress-related behaviour, suggesting that metabolic and ionic disturbances contribute to stress responses, as seen in other aquatic models [26].

### Proteomic Insights and Pathway Analysis

Proteomic profiling identified differentially expressed proteins across all treatment groups, with several linked to critical cellular functions, including cell survival, migration, and viability. The specific functions of differentially expressed proteins in zebrafish caudal fin tissue following ethanol exposure have not been extensively characterized. However, studies on zebrafish larvae have shown that ethanol metabolism leads to oxidative stress, which in turn induces endoplasmic reticulum (ER) stress and steatosis. These effects are mediated through the production of reactive oxygen species (ROS) during ethanol metabolism, suggesting that oxidative stress is a conserved aspect of alcohol-induced liver disease pathophysiology[27].

In the context of caudal fin regeneration, proteomic analyses have identified several proteins that are differentially regulated during the regenerative process. These include various keratins, cofilin 2, annexin A1, and structural proteins involved in maintaining cellular structure and architecture. Annexin A1, for instance, undergoes phosphorylation during regeneration, indicating its potential role in this process [28]. While direct studies on ethanol’s impact on protein expression during zebrafish fin regeneration are limited, the known effects of ethanol-induced oxidative stress and ER stress could potentially influence the expression and function of proteins involved in regenerative processes.

Glucose exposure influenced zebrafish caudal fin regeneration by modulating the organisms’ immune response, metabolism, and cytoskeletal dynamics. The upregulation of complement factor D precursor and serpin peptidase inhibitor also suggests enhanced immune activation, while LRP1 and clathrin interactor 1a indicate metabolic adaptations and increased vesicle trafficking, which could potentially aid in tissue repair. Additionally, the upregulation of dynamin-2 points to cytoskeletal reorganization, which may support the process of cell migration during regeneration. However, glucose exposure also led to the downregulation of glucosamine-6-phosphate isomerase 2andribosomal protein L22, indicating disruptions in protein synthesis and glucose metabolism. The suppression of peroxiredoxin-4 suggests increased oxidative stress susceptibility, while reduced catenin beta-1 expression may impair Wnt signaling, potentially affecting cell proliferation and tissue patterning. Overall, glucose exposure presents a dual effect such as enhancing immune and metabolic pathways while compromising oxidative balance and protein synthesis. Further studies are required to assess its long-term impact on regenerative efficiency and metabolic homeostasis.

Similarly, NaCl exposure altered zebrafish caudal fin regeneration by influencing key molecular pathways. The upregulation of fetuin-A, ARPC1A, SMC4, and tryptophan-tRNA ligase suggests enhanced cytoskeletal reorganization, chromatin stability, and metabolic adaptations to saline-induced stress, supporting cell proliferation and tissue remodeling during regeneration. Conversely, the downregulation of dipeptidyl peptidase 1, UDP-glucose 4-epimerase, ribosomal proteins, MHC class I UDA precursor indicates suppressed immune function, impaired protein synthesis, and disrupted lipid metabolism. These effects could hinder regenerative efficiency by reducing energy availability and slowing protein turnover. While NaCl exposure induces structural and metabolic adaptations, it simultaneously imposes constraints on immune response and protein synthesis, potentially affecting regeneration quality. Future investigations should focus on the long-term effects of saline stress on zebrafish physiology and tissue repair mechanisms.

The IPA analysis of canonical pathways and biological functions also provided insights into how different treatments affect regenerative and cellular processes in zebrafish. The observed patterns in pathway activation and disease-related functions suggest varied responses that align with existing literature on tissue injury, cellular stress, and immune responses.

The most significant pathways implicated include the GP6 signaling pathway, mitochondrial dysfunction, and the RHO GTPase cycle. The GP6 pathway is associated with the cellular response to injury and plays a role in tissue repair, while mitochondrial dysfunction points to potential oxidative stress within cells exposed to these treatments [29]. Downregulation in pathways like GP6 and actin-cytoskeleton signaling by 7dpa suggests that ethanol may initially stimulate immune responses that later taper off, compromising tissue repair and structural stability. The RHO GTPase pathway, known to regulate cell movement and adhesion, likely reflects the cellular reorganization needed for regeneration [30].

Interestingly, S Phase progression, deubiquitination, and TCR signaling were among the pathways associated with ethanol exposure, supporting prior findings that ethanol can disrupt cell cycle regulation and immune response pathways [31]. These pathways could explain the impaired regeneration and stressed behavior seen in the ethanol-treated group, as they collectively influence cellular stability, immune responses, and overall tissue viability.

The consistent upregulation of “organismal death” across all treated groups highlights a cellular stress response induced by the treatments. This may reflect increased metabolic burden and cell fate decisions or cellular apoptosis due to stressors [32]. Additionally, for the ethanol group, initial upregulation of activities such as cell migration, viability, and phagocytosis, followed by a downregulation at 7 dpa, suggests an early regenerative response that diminishes over time, potentially due to cumulative stress [25].

Ethanol exposure significantly upregulated pathways related to protein sorting and neutrophil extracellular trap formation initially, indicating that an early immune response may take place to counter tissue injury and promote cell sorting in regeneration. This is consistent with studies on ethanol’s effects on immune modulation and cytoskeletal dynamics in zebrafish, where early activation of immune pathways diminishes as cells undergo oxidative stress over time [33].

The glucose-treated group showed a consistent downregulation of metabolic pathways such as the S phase and RAF/MAP kinase cascade, indicating an inhibitory effect on cell cycle progression and proliferation. This may reflect cellular stress due to hyperglycemia, which can impede regenerative processes. Upregulated functions such as oxidative stress and apoptosis further confirm that increased glucose levels disrupt cellular homeostasis, likely affecting tissue recovery [15]. The suppression of pathways involved in Hedgehog signaling and ROBO receptors, which are critical for developmental processes and cell guidance, implies that glucose-induced metabolic disturbances inhibit pathways essential for tissue patterning and regeneration. The distinct upregulation of cell death-related activities across time points supports the idea of glucose’s role in inducing apoptosis through metabolic stress mechanisms [16].

NaCl treatment uniquely led to consistent downregulation in pathways related to fatty acid metabolism and immune functions, possibly due to ionic stress impacting cellular metabolism and immune responsiveness. This is corroborated by studies indicating that NaCl impairs lipid metabolism and immune pathways, contributing to tissue degeneration under stress [34]. Interestingly, pathways such as ROBO Slit signaling were upregulated initially, pointing to a brief activation of guidance-related mechanisms before systemic ionic imbalance. Downregulated immune and metabolic pathways by 7dpa suggest that continuous NaCl exposure may limit the regenerative capacity of tissues indicating that salinity impacts tissue repair. The NaCl group’s downregulation of KEAP1-NFE2L2 signaling, known for its role in oxidative stress responses, highlights a compromised antioxidant response, aligning with decreased cell viability and survival observed in this study.

### Implications for Regeneration Studies

Overall, our study highlights the distinct influences of ethanol, glucose, and NaCl on zebrafish regeneration, behaviour, and molecular pathways. Ethanol’s pronounced impact suggests that even low concentrations can inhibit regeneration and induce stress behaviours, likely due to its effects on oxidative stress and cell cycle regulation. In contrast, glucose and NaCl appeared to exert milder effects, though their impact on ionic balance and metabolic regulation remains significant. These findings contribute to a growing understanding of how small molecules modulate regeneration and stress responses in model organisms and may inform future studies on optimizing regeneration through precise modulation of small molecules. Additionally, the identification of key pathways offers potential therapeutic targets for enhancing tissue repair and addressing stress-induced impairment in regenerative models.

When comparing the effects of glucose, ethanol, and NaCl, it is evident that each molecule has a unique impact on the regeneration process, likely due to their distinct roles in cellular metabolism, signaling, and osmotic regulation. Our findings suggest a need for further investigation into the molecular mechanisms underlying these observations, exploring these small molecules in combination to understand potential synergistic or antagonistic effects on regeneration. By clarifying how these compounds influence cellular and molecular processes, we can better harness their potential for regenerative medicine applications, where controlled modulation of cell movement, viability, and migration could enhance healing outcomes.

## Conclusion

Our study underscores the complex interplay of small molecules in influencing regenerative mechanisms in zebrafish. The differential effects of glucose, ethanol, and NaCl point to the need for precise dosing and timing to harness these molecules for enhanced tissue repair. This study provides evidence that each treatment alters zebrafish regenerative pathways differently, reflecting unique molecular responses to chemical-induced stress. These findings contribute to our understanding of chemical resilience and toxicity in regenerative biology, reinforcing zebrafish as a suitable model for studying tissue repair mechanisms in response to varied small molecule interventions. This study also highlights the potential pathways of small molecules through various tissues and cells which prove to play an important role in the designing of therapeutic therapeutic or precision medication drugs.

## Abbreviations

hpa: hours post amputation
dpa: days post amputation

## Declaration Statement

## Ethics approval

The experimental protocol was approved by Centre for Cellular and Molecular Biology institutional animal ethics committee (IAEC/CCMB/Protocol #50/2013).

## Consent for Publication

Not Applicable

## Data Availability

All the data generated in the study were presented in the manuscript and available in PRIDE database. Project Name: Effect of small molecules in zebrafish caudal fin regeneration. Project accession: PXD060144

## Authors’ contribution

APV: Performed experiment, analysed data and wrote manuscript.

ST: Performed experiment.

CP: Performed experiment

AA: Wrote the manuscript.

II: Wrote the manuscript.

MI: Conceived the idea, wrote the manuscript.

All authors read and approved the final manuscript.

## Acknowledgements

This work was supported by CSIR. The authors are thankful to Ms. Noorul Fowzia for critically reviewing the manuscript.

## Supplemental Data

This article contains supplemental data.

